# Plant Scaffolds Support Motor Recovery and Regeneration in Rats after Traumatic Spinal Cord Injury

**DOI:** 10.1101/2020.10.21.347807

**Authors:** Daniel J. Modulevsky, Charles M. Cuerrier, Maxime Leblanc-Latour, Ryan J. Hickey, Ras-Jeevan K. Obhi, Isabel Shore, Ahmad Galuta, Krystal L. A. Walker, Eve C. Tsai, Andrew E. Pelling

## Abstract

As of yet, no standard of care incorporates biomaterials to treat traumatic spinal cord injury (SCI). However, intense development of biomaterials for treating SCI has focused on fabricating microscale channels to support the regrowth of axons while minimizing scar formation. We previously demonstrated that plant tissues could be decellularized and processed to form sterile, biocompatible and implantable biomaterials that support cell infiltration and vascularization *in vivo*. Vascularized plant tissues contain continuous microscale channels with geometries relevant for supporting neural regeneration. We hypothesized that decellularized vascular bundles would support neural regeneration and motor recovery in SCI. Sprague Dawley rats received a complete T8-T9 spinal cord transection and were implanted with acellular plant-derived scaffolds and allowed to recover over 28 weeks. Animals that received the scaffolds alone, with no other therapeutic compounds, demonstrated a significant and stable partial improvement in motor function compared to control animals as early as week 4 post-injury. Hind-limb motor function did not deteriorate over the remaining 28 weeks. Histological analysis revealed minimal astrocyte scarring at the spinal cord - scaffold interface, aligned axonal projection through the scaffolds, populations of serotonergic neurons and Schwann cells, laminin and collagen deposition and the presence of blood vessels. Axonal reconnection via the scaffold was also confirmed by Fluro-gold retrograde tracing. Taken together, our work defines a novel route for building upon naturally occurring plant microarchitectures to support the repair of the spinal cord post-injury. Notably, these results were achieved without the use of growth factors, stem/progenitor cells, or any other interventions.

## INTRODUCTION

The incidence of SCI is 54 per million in developed countries, with the leading causes being traffic accidents, falls, sports-related injuries, and neoplasia. Due to lack of effective treatments, life expectancy with SCI have not improved since the 1980s and are significantly lower than that of persons without SCI^1,2^. Although treating the devastating loss of motor function is the ultimate clinical goal for patients, recovery of bowel, bladder, sexual and tactile function can significantly improve patient quality of life^3^. While there are no accepted therapies to treat the underlying issue of astrocyte scar and or necrotic cyst formation at the epicenter of the injury^4–8^. Researchers and clinicians have been developing biomaterials that can promote axonal regrowth, sequesters scar tissue and support new blood vessel formation into the biomaterial, which ultimately aids in the recovery of function after spinal cord injury^9^. There has been an intense effort to create scaffolds with 3D architectures designed to achieve this goal utilizing all manner of microfabrication techniques^10–14^. In many previous studies, biomaterials for SCI require supplementation with other factors such as neural progenitor cells (NPCs), pharmaceuticals, or growth factors (alone or in combination) to achieve a significant desired therapeutic effect^12,13^ Moreover, tissue regeneration and improvement in motor recovery are only possible with biomaterials and combined strategies^12,15–19^.

Recent advancements in microfabrication techniques has allowed for the development of novel SCI biomaterials, such as those with internal channeled architectures that mimic the underlying architecture of the spinal cord in a rat model^12^. In this previous work, the channels led to the sequestration of astrogliosis and appeared to support axonal projection. However, motor recovery of the hind limbs was only evident when the scaffolds were loaded with NPCs^12^. The performance of many natural and synthetic polymers has been investigated in several SCI animal models^6,10– 13,20–30^. In a previous rat study, animals that received scaffolds filled with methylcellulose hydrogels demonstrated the greatest tissue regeneration and motor recovery compared to other candidate biomaterials^27^. A review of these studies, reveals that natural rather than synthetic polymers, specifically cellulose, tend to avert severe foreign body responses (FBR) and astrogliosis in SCI^27,31–35^.

Previous work from our group and others has shown that the cellulosic microarchitecture in decellularized plant tissues is biocompatible *in vitro*^36–44^ and *in vivo*^37,40,45^ and appear to have many potential applications in tissue engineering and regenerative medicine^25,26,50,27,32,39,40,46–49^. Cellulose is the building block of the cell wall of plants and is an organic polysaccharide of linear syndiotactic D-glucose molecules linked through β(1→4) glycosidic bonds. Most vertebrates do not directly express the cellulase enzymes required to break down the β(1→4) glycosidic bonds, and as such, cellulose provides very little nutritional energy ^51^.

After implantation, plant-derived decellularized scaffolds incite low foreign body response (FBR) levels in the surrounding epidermis tissue^37,40^. Furthermore, fibroblasts are observed migrating from the surrounding epidermis throughout the scaffold, depositing a new collagen extracellular matrix. In conjunction, blood vessels are regularly observed throughout the scaffold supporting the migrating cells^37,40^. The utility of such plant-derived scaffolds for regenerating tissues in animal models has now been demonstrated in soft tissues ^37,40^ and bone^45^.

Here, we were inspired by the observation of dense linearly arranged-parallel microchannels forming vascular bundles (VBs) in the stalks of *Asparagus officinalis* which possess an architecture with striking similarities to scaffolds designed to repair SCI in animal models^27,31,52– 57^. Tracheophyte plants, including clubmosses, horsetails, ferns, gymnosperms and angiosperms, have evolved distinct vascular structures that support the transport of water and nutrients via the xylem and phloem^58^. The xylem is lignified channels specialized for water transport and nutrient, derived from the root tissue, and function as a support function in the stem of plants: phloem structures, transport products of photosynthesis from source tissues to sink root tissues^58^

This study demonstrates that scaffolds composed of decellularized *Asparagus officinalis* possess structural features that support neural tissue regeneration and motor recovery in rats with a complete T8-T9 spinal cord transection injury. Animals that received plant-derived scaffolds demonstrated improved motor function as early as 4-week post-injury and remained stable throughout the remainder of the six-month study. Results reveal that animals with scaffolds had higher NF200 positive regenerated axon densities through the scaffold and surrounding host tissue. The functionality of the projecting axons was also investigated with retrograde tracer studies, which revealed an axonal reconnection through the implanted scaffolds. Additionally, histological analyses reveal that the larger diameter vascular bundles in the implanted scaffolds act to sequester astrocyte-rich scar tissue, contributing positively to tissue regeneration and motor recovery. Consistent with our previous studies, the plant-derived scaffolds remained stable over time and became vascularized along their length^37,40^. Importantly, this work demonstrates that the observed motor recovery was due to the presence of the scaffolds alone, with no therapeutic stem cells or other interventions required in contrast to numerous similar studies. Taken together, our work defines a novel route for exploiting naturally occurring plant microarchitectures as platforms to support the repair of functional spinal cord tissue.

## 2. RESULTS

### 2.1 Physical properties of the Vascular Bundles for SCI scaffolds

In the plant-derived scaffolds studied here, VBs are circularly arranged and separated by parenchyma tissue with an average spacing of 612±70μm (n=11). VBs aid in the efficient transport of water, nutrients and biomolecules throughout the plant. Notably, the cell wall structure of VBs is preserved after decellularization (Fig. 1a, b). Scanning electron microscopy (SEM) of the VBs reveals a variety of tissue structures with characteristic diameters, such as xylem channels (51±15 μm, n=11), sieve tubes (40±16μm, n=11), parenchyma (35±8μm, n=8) and the phloem (9±2μm, n=11) (Fig. 1c-e, Supplementary Fig. 1). Due to the distribution of microchannels in their architecture, we hypothesized that decellularized vascular plant tissues could be exploited as lignocellulosic scaffolds to repair complete spinal cord injury in a rat model. On average, each scaffold contained 11±2 VBs (n=4), each composed of 35±5 continuous microchannels (n=11) (Fig 1 a,b). These channels were consistent in diameter and orientation throughout the entire scaffold and emerge in the same position on either end (Supplementary Fig. 2). Scaffolds were selected that contained the maximum number of VBs to promote invasion of regenerating axon projections in a completely transected spinal cord. The scaffolds are mechanically anisotropic due to the linear orientation of the VBs along the plant stem with an elastic modulus of 148±53kPa (n=10) or 12±4 kPa (n=10) when measured parallel or perpendicular to the long axis, which is within the range of healthy rat spinal cords^59^ (Supplementary Fig. 3).

**Fig. 1.**
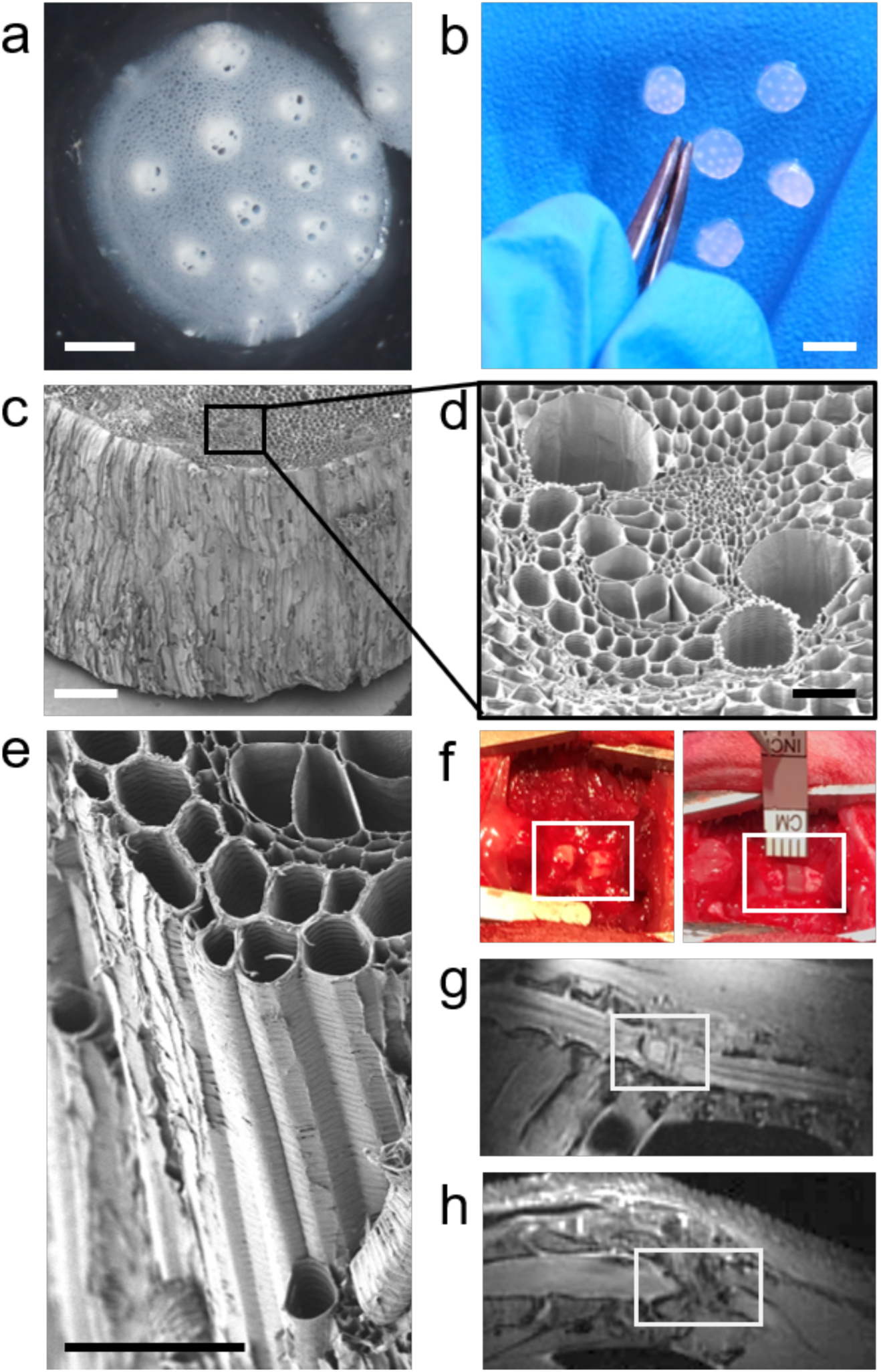
Plant-derived scaffolds for spinal cord injury. **a)** Decellularized plant-derived scaffold with visible VBs surrounded by parenchyma (bar=1mm). **b)** VBs are even visible to the naked eye (bar=4mm). **c)** SEM reveals that scaffolds also have a grooved outer topography (bar=300μm). **d)** A cross-section at higher magnification of an individual VB reveals a highly channeled microarchitecture (bar=100μm). **e)** VB channel microarchitecture runs the entire scaffold length (bar=100μm). **f)** An exposed fully transected spinal cord (left, box) and a scaffold after implantation but prior to fibrin deposition (right, box). **g**) Sagittal MRI of T8-T9 injury (box) containing a scaffold after 4 weeks, and **h)** Continuing degeneration of the surrounding spinal cord in animals not receiving the scaffold after 4 weeks.

### 2.2 Complete Spinal cord Injury and Scaffold Implantation

In animals with a complete T8-T9 spinal cord transection, the scaffold group (n=23 animals) had grafts inserted with lengths to match the gap distance of the stumps (Fig 1f). The scaffolds were placed with the long axis of the VBs parallel to the spinal cord. Fibrin was applied across the dorsal surface of the scaffold to fix it between the stumps. The control group (n=11 animals) did not receive the scaffold, and the fibrin was applied to the gap formed between the stumps. After 4 weeks, animals were imaged with magnetic resonance imaging (MRI) to confirm that the scaffold remained in place and was not damaged (Fig. 1g). The rostral and caudal stumps of the transected spinal cord stay aligned and in contact with the scaffold. In contrast, typical symptoms of progressing Wallerian degeneration and the formation of cyst and syrinx typical to intermediate and chronic secondary spinal cord injuries were apparent in the control animals ^3,12,24,25,60–62^ (Fig. 1h, Supplementary Fig. 4).

### 2.3 Hind limb Motor Recovery

The motor recovery of the rats was blindly assessed using the established Basso, Beattie and Bresnahan (BBB) locomotor scale^63^ (n-values for each time point found in Supplemental Table 1). Rats that received the scaffold had a statistically significant functional recovery starting at week 4 (p=0.030655) (Fig 2a). Though both groups were observed to display some motor recovery, the degree of recovery was significantly higher in animals that received the scaffold. By week 4, the motor recovery of control animals plateaued at 2.3±0.5 (extensive movement of one joint). Conversely, animals that received the scaffold displayed continued improvement until week 7, after which the BBB score plateaued at 5.5±0.1 (slight movement of the 2 hindlimb joints and extensive movement of the third, p=2.4×10^−7^). A small minority of animals that received the scaffold (n=3) demonstrated instances of BBB scores between 7 to 9. These particular animals were placed on a DigiGait treadmill and carefully encouraged to gait. The selected animals demonstrated limited weight-supported plantar stepping and coordination (Supplementary Fig. 4). Overall, the motor skills of all animals receiving the scaffold eventually plateaued and did not decrease significantly over the remainder of the entire 28 week study (Fig. 2a). As well, no animals were removed from BBB analysis due to poor motor recovery and the data in Fig. 2a is an average of all data from all study animals. To rule out reflex adaptation of the rhythmogenic elements, animals at their respective endpoints were selected at random for re-transection surgery at the T13 vertebra^43^. Animals were allowed to recover for one-week prior motor skill re-assessment with the BBB locomotor scale. In the scaffold and control groups, all motor recovery was lost and animals demonstrated BBB scores of 0.3±0.2 and 0.1±0.1 at week 14, respectively (p=0.251289794) (Fig. 2a) which we consistent with the retransected week 28 animals.

**Fig 2.**
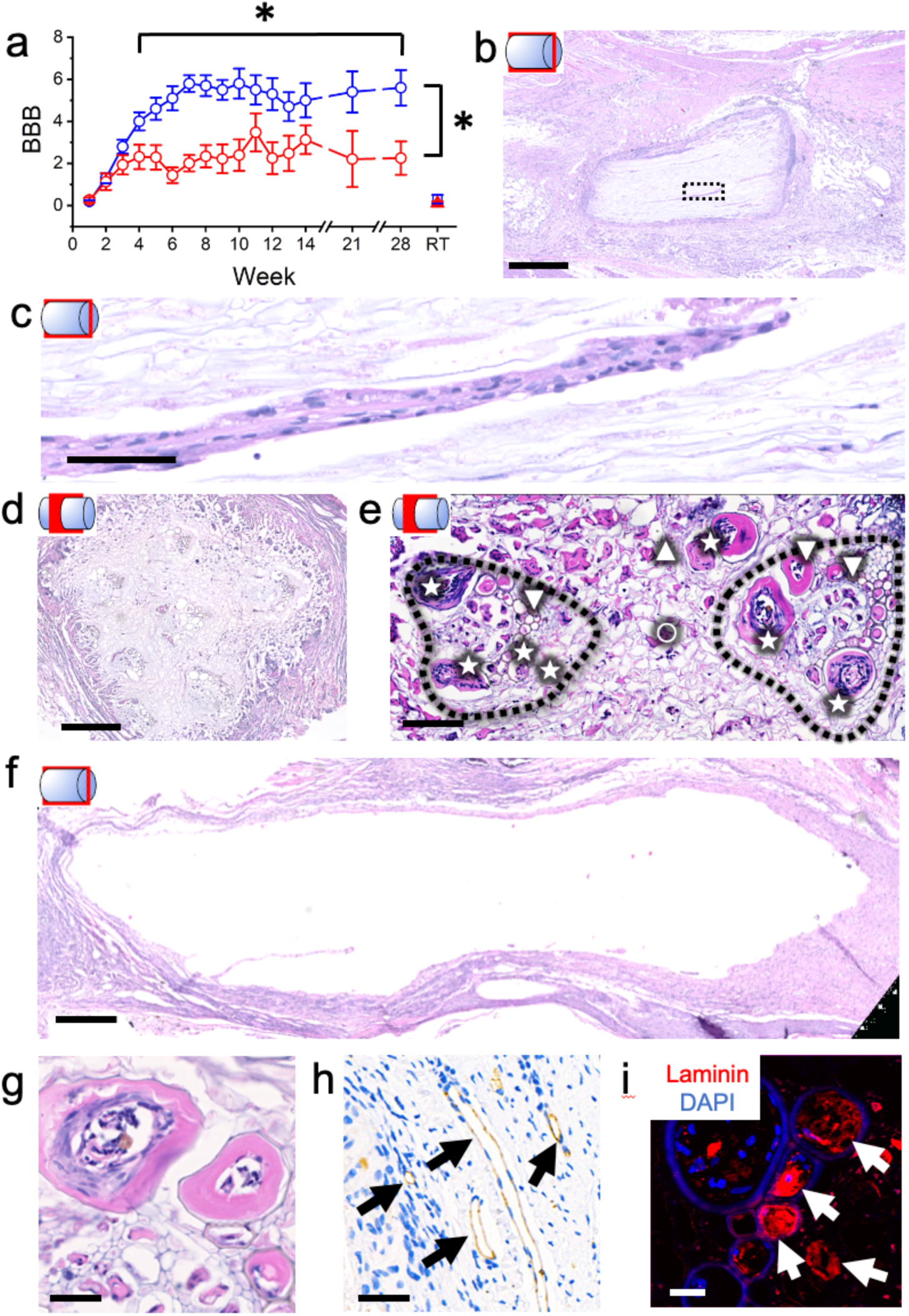
Motor recovery and histology after 14 weeks post-injury. **a)** BBB motor scores after complete transection in animals with (blue) and without a scaffold (red) are significantly different from week 4 and onwards. Re-transection (RT) at week 14 causes the scores of the scaffold and control groups to return to 0.3±0.2 and 0.1±0.1, respectively. **b)** Sagittal H&E of the scaffold. The scaffold is observed integrated within the initial injury site of the spine. Host tissue is observed surrounding the scaffold and within the VBs and the parenchyma even at the core of the scaffold (bar=1mm). **c)** A magnified view of the ROI in b demonstrating host cell invasion along a VB (bar=400μm). **d)** Wide field axial H&E through the scaffold. VBs are infiltrated with host cells (bar=400μm). **e)** H&E of two VBs (dotted lines). Blood vessels (∇)and granulation tissue (★) are observed in the VBs. The parenchyma (o) between the VBs contains fibroblasts (Δ) (bar=100μm) **f)** Sagittal H&E of a control animal spinal cord for comparison (bar=500μm). Blood vessels with thick endothelial linings can be observed in the VBs and parenchyma as confirmed with **g)** H&E (bar=50μm), **h)** CD31 with a DAB chromogen and hematoxylin counterstain. Arrows indicate Blood vessels (bar=25μm) and **i)** laminin immunofluorescence staining (bar=20μm).

### 2.4 Histology of SCI scaffolds post-implantation

Animals were euthanized and tissues were harvested at the respective endpoints of 14 (n=16) and 28 weeks (n=18). Tissues were then sectioned in either an axial or sagittal orientation (Fig. 2). During the dissection, the scaffold was found to be well integrated to the two stumps of the spinal cord and could support the weight of the CNS tissue when carefully dissected out (Supplementary Fig. 5). These observations are consistent with MRI, which revealed that the scaffold formed a physical bridge between the spinal cord stumps (Fig. 1g, h). The control groups were difficult to dissect as the stumps loosely attached to one another (Supplementary Fig. 5). At 14 weeks, hematoxylin and eosin (H&E) staining reveals that the scaffold retained its pre-implant size and the parenchyma and VBs were infiltrated with surrounding host cells (Fig. 2b-d, Supplementary Fig. 6-8). Host cells could be observed deep within the VBs (Fig. 2c, e) as well as projecting into the VBs at the interface between the scaffold and spinal cord (Supplementary Fig. 6). Host cells could also be observed within the interstitial spaces formed by the overlapping plant cell walls in the parenchyma tissue (Fig. 2e). Although macrophages were present, there was no observable degradation of the scaffold. As well, almost no foreign multinucleated cells associated with chronic FBR, basophils and lymphocytes were observed (Fig. 2e). Notably, at 28 weeks, H&E imaging revealed results highly consistent with the 14 week time point (Supplementary Fig. 7). The control animal injury site was void of cell infiltration, with the formation of large cysts resulting in a greater distance between the stumps as is typical of pathologies related to SCI injuries^3^ (Fig. 2f). Blood vessels were also observed throughout the scaffold as identified with H&E and were found within 86% of all the VBs (n=10) leading to vascularization through the injury site (Fig. 2g). Blood vessels were further confirmed with CD31 (n=4) and laminin (n=3) staining (Fig. 2h, i). Blood vessel formation within decellularized plant scaffolds is highly consistent with all of our previous *in vivo* studies^37,40^.

Laminin (Fig. 2i, Supplementary Fig. 9) and collagen (n=6, Supplementary Fig. 10) were also observed deposited throughout the scaffold, highlighting the innate ability of the scaffold to support host tissue integration and remodeling. The most prominent channels of VBs were observed to incorporate a large amount of scar tissue and appear to contain a significant amount of collagen deposition along the length of the scaffold (Fig. 2e, Supplementary Fig 9, 10).

### 2.5 Host cell invasion into the SCI scaffold post-implantation

Astrocyte scarring was identified by staining for glial fibrillary acidic protein (GFAP) (n=13). The sagittal section of rostral and caudal ends of the scaffold-spinal cord interface displayed low GFAP scarring compared to the control animal cord stumps (Fig. 3a-c). The rostral/caudal scaffold can be seen well incorporated into the cord with limited scarring seen at each respective interface after 14 weeks (Fig. 3a, b). As seen in the H/E, control animals had significant degeneration in both stumps (Fig. 2f) The ischemic penumbra has astrocyte scarring typical with such a significant SCI. (Fig 3 c, d). At 14 weeks, the normalized area (Supplementary Fig. 11) of GFAP labelled astrocyte scarring (normalized to total tissue area) was quantified at the rostral and caudal interfaces of the biomaterial (n=8) with the spinal cord and was found to be 0.08 ± 0.02. This is significantly less than the normalized GFAP area in 14-week controls (n=3) in a similar region which was found to be 0.80 ± 0.05, p=4.1719 ×10^−17^) (Fig. 3e). These results are consistent with the limited astrocyte scarring in previously reported synthetic cellulose-based scaffolds^6^. Notably, when GFAP-positive astrocytes were observed in the scaffold and surrounding tissue, they were generally located within the VBs (Supplementary Fig. 12) (n=5). Though the precise role of glial scar tissue is still unknown, scarring was observed to be segregated within the large diameter channels rather than the parenchyma, consistent with the H&E and Masson Trichrome (n=3) staining at 14 and 28 weeks (Supplementary Fig. 6-8 and 10). The presence of both collagen and laminin deposition within the scaffold was also observed (Supplementary Figure 9 and Supplementary Figure 10). Collagen tended to also be sequestered within the channels and associated with GFAP signal and glial scarring, while laminin deposition was observed both within the channels and surrounding parenchyma.

**Fig 3.**
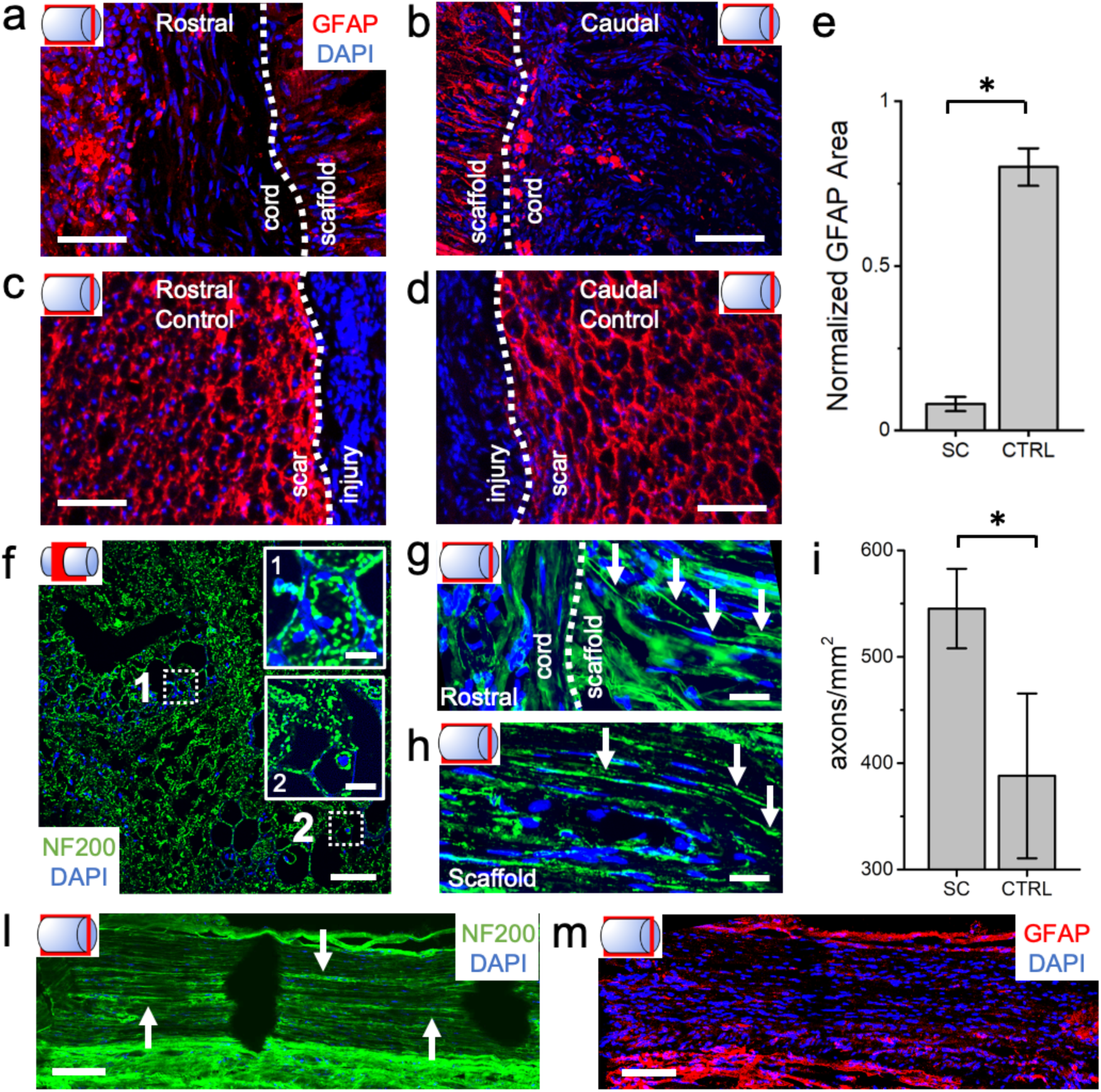
NF200 and GFAP staining 14 weeks post-injury. **a)** Rostral end, sagittal view of the scaffold interface (dotted line) with spinal cord stained for GFAP (red) and DAPI (blue) (bar=100μm, for a to d). **b)** Caudal end, sagittal view of the scaffold-cord interface **c)** Rostral end, sagittal view of the stump formed from the initial injury. GFAP staining is primarily segregated into the scaffold itself rather than in the surrounding spinal cord tissue. **d)** Caudal end, sagittal view of the stump formed from the initial injury. **c)** Rostral end, sagittal view of the stump formed from the initial injury. In the control animals a significant amount of GFAP staining is observed in the cord tissue in contrast to the animals which received scaffolds. **e**) The normalized area of GFAP labelled astrocyte scarring at the rostral and caudal interfaces of the biomaterial or injury and the spinal cord in animals that received scaffolds (SC) or were controls (CTRL). This data quantifies the observations that the surrounding spinal cord tissue contains significantly less GFAP positive cells than animals which received scaffolds. **f)** Axial section within the rostral end of the scaffold stained for NF200 (green) and DAPI (blue) after 14 weeks (bar=100μm) reveals punctate structures consistent with the morphology of axon cross sections. Magnification of the boxed areas **1** and **2** are shown in the inserts (bars=20μm). **g)** Sagittal view of the scaffold-cord interface stained with NF200 (green) and DAPI (blue) after 14 weeks (bar=25μm, for g and h). The cord-scaffold interface is marked with the dotted line and several NF200 filaments of axons are observed entering the scaffold (arrows). **h)** Sagittal view within the scaffold reveals NF200 labelled axons within the midpoint the scaffold. **i)**The density of NF200 labelled axons within the scaffold and within the injury site of control animals. **l)** Sagittal view of the dorsal exterior surface of the scaffold. Axons appear in a linear orientation (arrows) (bar=125μm). **m)** Sequential sagittal slides stained with GFAP (red) and DAPI (blue) reveals little scarring in tissues on the dorsal surface of the scaffold consistent with observations at the rostral and caudal ends (bar=125μm).

Host axons, labelled with neurofilament protein NF200 (n=18), were observed regenerating readily throughout the entire scaffold in a linear orientation along the median axis of the spinal cord (Fig. 3f). Axons were observed extending from the spinal cord into the scaffold at both ends, as well as at its midpoint, and were generally oriented parallel to the long axis of the implant (Fig 3g, h). Axon density (545±37 axons/mm^2^, n=12) inside the 1.27 mm^2^ cross sectional area scaffolds was significantly higher (p=0.01859) than in the tissue of the control scar tissue (368±62 axons/mm^2^, n=8) (Fig. 3i and Supplementary Fig. 13). In control animals, axons were only observed in limited sections of the scar tissue of the stumps. As well, the regenerated tissue on the surface of implants contained axons that were observed to align with exterior linear topography of the scaffold, consistent with contact guidance^64^ (Fig 3l). Inhibited scar tissue formation (low GFAP signal) was also observed in these regions as well (Fig. 3m). These results suggest that the scaffold provides both an exterior and internal microenvironment to promote the growth of native axons, while inhibiting astrogliosis.

We also stained for 5HT at 14 (n=6) and 28 (n=6) weeks post-injury to investigate the presence of serotonergic neurons during regeneration. As expected, 5HT positive axons were present in the spinal cord tissue rostral to the injury in control animal and in animals receiving the scaffold (Supplementary Figure 14)^65^. However, although there was some indication of 5HT-positive structures within the scaffold and in the spinal cord caudal to the injury, the density and signal quality was extremely low making identification difficult.

### 2.6 Retrograde tracing of axonal projections within the Scaffold

To determine the functionality of axons projecting into the scaffold from the rostral and caudal ends, we utilized retrograde axonal tracing with Fluorogold (FG). At 14-weeks post-injury, animals with scaffolds (n=8), and controls (n=4), underwent a second complete transection at thoracic-13 vertebrae (Fig. 4a) and had FG applied to the rostral aspect of the cut cord to label the motor neurons in retrograde fashion. FG labelled axons were observed inside the scaffold at both the caudal (proximal to the loading site) and rostral (across the scaffold) scaffold interfaces with the spine (Fig. 4b, c). In animals receiving the scaffold, rostral sections of the spinal cord contained neuronal cell bodies with punctate structures resembling FG loaded vesicles (Fig. 4d). As well, many DAPI-labelled nuclei did not display any correlation to FG labelled cell bodies demonstrating the specificity of the FG retrograde tracer (n=4). Some samples also had the scaffold sectioned and counterstained with NF200 (Fig. 4e). Sagittal sections revealed FG signal and cell bodies correlating with presence of NF200 positive axonal structures deep within the scaffold (n=4). We observed a low FG signal in control experiments (n=3) and a lack of nonspecific FG wicking through the scaffold in negative controls (Supplementary Fig. 15). These results confirmed that FG-positive cells were a result of retrograde axonal transport. FG intensity and FG positive cell counts were quantified in the spinal cord rostral (n=4) and caudal (n=4) to the scaffold (Fig. 4f). The number of FG-positive cells on the rostral side of the injury was significantly decreased compared to the caudal end in control animals (p=0.01102), which was not the case in animals that received a scaffold (p=0.77785). Likewise, FG intensity nearly disappeared across the transection in the control animals (p=0.00278). Conversely, in animals that received a scaffold, FG intensity remained constant (p=0.59004). The results demonstrate the presence of axonal connections between neurons on the caudal side of the scaffold to axons and cell bodies on the rostral side (Supplementary Fig. 16).

**Fig 4.**
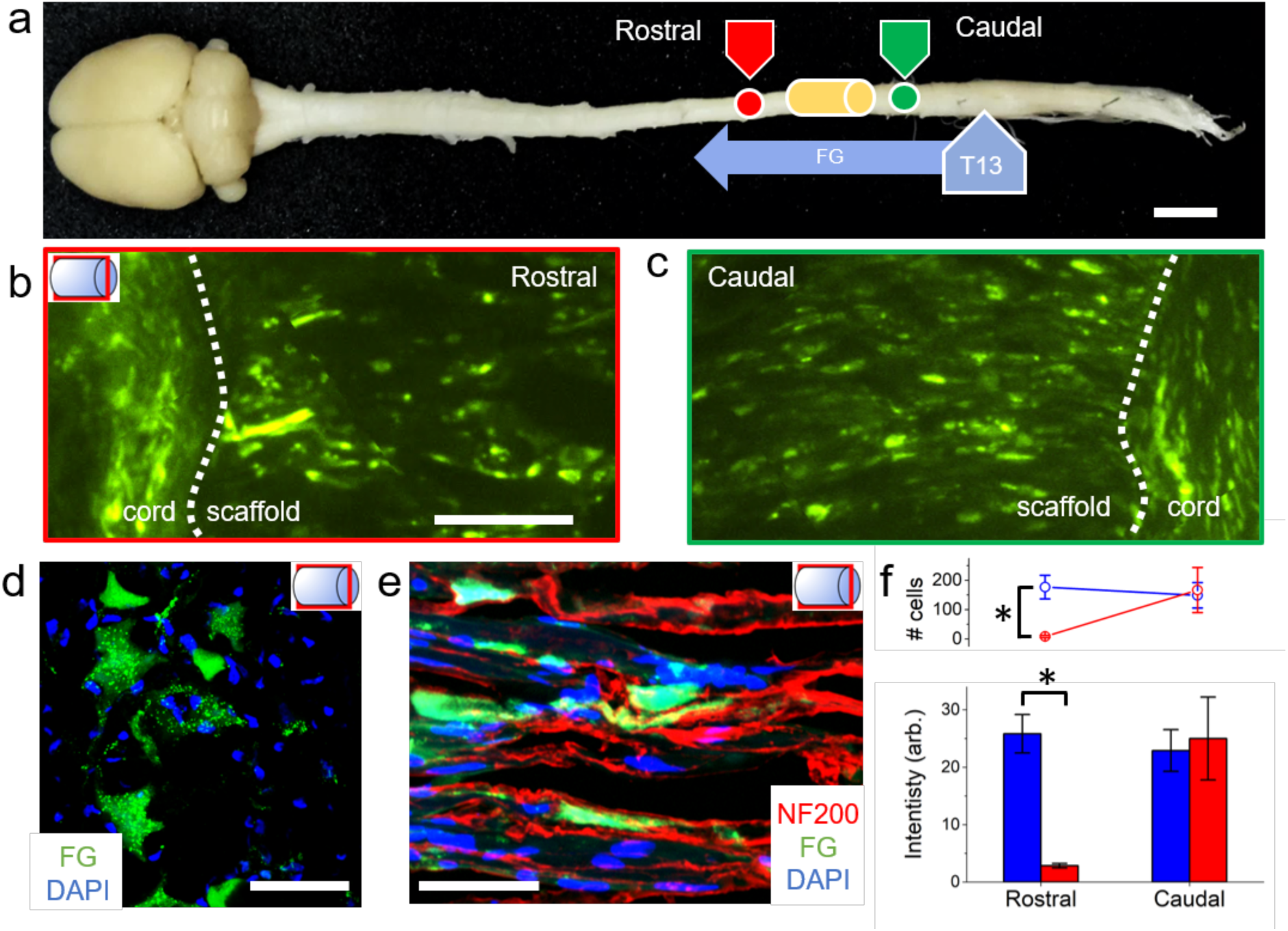
Retrograde axonal tracing 14 weeks post-injury. **a)** Animals received a secondary transection at T13 caudal to the original injury, and FG, a retrograde axonal tracer was loaded into the new site (healthy CNS shown for illustration, scaffold shown as a yellow cartoon, FG direction of transport shown as a purple arrow. One week later, sagittal sections of the **b)** rostral (red position in **a**) and **c)** caudal (green position in **a**) ends to the original site of injury or scaffold reveal projecting axons at the scaffold-tissue interface (dotted line) (bar=100μm for **b** and **c**). The scaffold interface has FG positive axons extending from the scaffold into the host tissue at both ends. **d)** Axial section of the tissue rostral to the scaffold (red position in **a**) double labelled with FG and DAPI (bar=50μm). FG vesicles within the cell bodies are present in several cells. Many nuclei are observed throughout the tissues without any correlation to FG, demonstrating the specificity the tracer (bar=50μm). **e)** A sagittal magnified view of the scaffold labelled with NF200 (red), FG (green) and DAPI (blue). FG signal correlates with NF200 signal in axons within the scaffold (bar=50μm). **f)** The number of FG positive cells and corresponding FG intensity significantly decreases after the injury site in controls (red) (p=0.01102 and p=0.00278, respectively) as opposed to animals with scaffolds (blue) (p>0.05).

Finally, in addition to NF200, FG and 5HT staining, we also performed S100 staining of the scaffold at 14 and 28 weeks (n=3). The results confirmed the presence of Schwann cells in the scaffold (Supplementary Fig. 17). Furthermore, animals receiving FG demonstrated the presence of some, but not all, axons potentially ensheathed by Schwann cells or remyelinated host axons. Interestingly, the S100 signal was mainly observed within the parenchyma rather than the VB channels.

## 3. DISCUSSION

There has been significant hope in treating SCI with the development of a multitude of natural and synthetic biomaterials^25–27,32,39,46–49^. Scaffolds composed of chitosan, collagen, hyaluronic acid, silk, methacrylate-derived poly-(2-hydroxyethyl) methacrylate and polyethylene glycol (PEG) have all demonstrated promising potential for SCI repair. Recently, researchers were able to bio mimic complex CNS structures in 3D printed PEG scaffolds^12^. This extremely innovative approach led to the creation of scaffolds with microscale channels that mimicked the underlying architecture of the spinal cord in a rat model^12^. However, only once loaded with NPCs did these scaffolds result in animals achieving a BBB score of 6.6±0.5 after a complete transection injury. Consistent with that study, the channels in our scaffolds aided the sequestration of astrogliosis, allowing for axonal projection through scaffold parenchyma and VBs^7^. Several previous studies have also demonstrated significantly increased BBB scores, but only after scaffolds were loaded with NPCs and/or complementary agents to aid in neuroregeneration^12,16,17,49,66–68^. A common characteristic of these studies is that NPCs and other therapeutic approaches are clearly required to stimulate any significant motor function recovery. In contrast, we observed improved motor function similar to previous studies, attributed to only the acellular plant scaffolds. Furthermore, to our knowledge, only a small minority of studies have demonstrated that a non-degradable stable scaffold can lead to a consistent improvement in motor function over 28 weeks in a rat model^15,69^. Here, we have shown that naturally derived plant cellulose scaffolds are biocompatible and become vascularized after implantation^37,40^. The low FBR and observed vascularization in our scaffolds are likely to have aided both tissue infiltration/regeneration and functional recovery without the need for therapeutic factors.

Biodegradability is often thought to be necessary for implantable biomaterials^28^. However, natural biomaterials such as chitosan and cellulose have shown some success in SCI while being non-degradable^27,32^. Plant-derived cellulose scaffolds are non-resorbable; however, they are highly biocompatible after implantation, structurally stable, and support tissue integration^37,40^. The scaffolds maintained their physical dimensions and did not demonstrate any native degradation throughout the 6 months of implantation. Importantly, the scaffolds did not demonstrate inhibitive symptoms of chronic biomaterial FBR. H&E, CD31 and laminin staining consistently revealed a clear distribution of blood vessels in the scaffold. At 14 weeks, 86% of all VBs channels were vascularized to some degree. The formation of new blood vessels in SCI has been attributed to improved functional recovery following injury in animal and human models following stroke^70^. Taken together, these results demonstrate the beneficial properties of acellular plant vascularized scaffolds to promote tissue regeneration and motor function recovery after SCI in rodents.

Furthermore, the long-term stability of the scaffold mitigates the risk of surgical removal. Host cells were able to infiltrate and pass through the entire length of the VBs. As well, it is important to highlight that the scaffold supported the projection of a significant number of NF200 positive axons across the injury site. Such axons were only observed when the scaffold was present and were not observed in the injury site of control animals. Conversely, serotonergic 5HT-labeled axons were also observed in the cord tissue rostral to the injury site. However very few 5HT-positive structures were observed in the scaffold or caudal to the injury. Importantly however, scar astrocyte tissue was significantly decreased at the biomaterial/spinal cord interface, and any GFAP positive cells within the scaffold appeared to be largely sequestered in the VBs. We speculate that this phenomenon may be beneficial for the observed recovery, however much more research will be required to investigate the significance of this observation. Finally, FG results demonstrated that plant-derived scaffolds support axonal reconnection from the stumps across the injury site. Indeed, some FG-positive axons were also observed to be colocalized with S100. Future research will be conducted to investigate and characterize the myriad of axon types that are supported by the scaffold and delve deeper into the potential mechanisms underlying the observed recovery in this study.

The primary objective of this study was to establish that the microarchitectures found in plant-derived scaffolds can be exploited to repair neuronal tissues in SCI and offer a potential platform for future discovery and innovation. These results demonstrate that such scaffolds can support regeneration of functional neural tissues in the most extreme model of traumatic SCI at levels consistent with previous studies^12,16,27^. Notably, these results were achieved without additional pharmaceuticals or therapeutic cells. However, such combinatorial approaches can be explored in the future to investigate synergistic effects leading to improved regeneration and motor skill recovery beyond the levels observed here. Acellular lignocellulosic scaffolds can be seeded with a vast array of cell types and functionalized to include extracellular matrix proteins or neural growth factors through hydrogel loading and/or surface functionalization^20,71–74^. The emergence of plant-derived scaffolds for tissue engineering has opened up many new possibilities to regenerate target tissues of interest, including soft tissues, muscle, and bone, by exploiting plant microarchitectures^36–40,44,45^. The results presented here demonstrate that such approaches can be exploited to aid in the regeneration of complete SCI, an incredibly complex injury model with low endogenous repair. The results point to exciting potential patient-treatment strategies in which plant-derived scaffolds might be deployed in combination with other therapeutics.

## 4. MATERIAL AND METHODS

### 4.1 Biomaterial Production

This protocol is based on our previously published works^9–11^. Asparagus (*Asparagus officinalis)* was purchased from local supermarkets. The asparagus was stored at 4°C in the dark for a maximum of one week and kept hydrated. To prepare the scaffolds, the asparagus stalks (with a diameter 14–17 mm) were washed, and the stalks’ end was cut to remove any dried tissue. The scaffolds were cut at different lengths using a #820 microtome (American Optical Company). The thickness of the asparagus scaffold was adjusted with the z-position block. The desired length of asparagus was cut with microtome blades (Westin Instruments Boston) in a quick motion to create two perpendicular surfaces with a precise length ranging from 0.2 mm – 1.6 cm. The resulting sections were then measured with a Vernier caliper. A 4 mm biopsy punch was used to cut out cylindrical sections close to the edges of the tissue to maximize the number of VBs. Effort was made to avoid the central fibrous tissues common in all angiosperm plants. Asparagus samples were placed in sterilized 2.5ml microcentrifuge tubes, and 2ml of 0.1% sodium dodecyl sulphate (SDS) (Sigma-Aldrich) solution was added to each tube. Samples were shaken for 48 hours at 180 RPM at room temperature. The resulting cellulose scaffolds were then transferred into new sterile microcentrifuge tubes, washed and incubated for 12 hours in phosphate-buffered saline (PBS) (Sigma-Aldrich). Following the PBS washing steps, the asparagus were then incubated in 100 mM CaCl2 for 24 hours at room temperature to facilitate the removal of any of the remaining SDS. The samples were washed 3 times with dH2O and then sterilized in 70% ethanol overnight. Finally, they were then washed 12 times with sterile saline solution and stored in saline. At this point, the samples were immediately used or stored at 4°C for no more than 24 hours.

### 4.2 Young’s Modulus Testing

Scaffolds were loaded onto a CellScale UniVert (CellScale) compression platform. The Young’s modulus was measured by compressing the material to a maximum of 10% strain at a compression speed of 50 µm/s. The force-indentation curves were converted to stress-strain curves and fitted in Origin 8.5. The Young’s modulus was extracted from the elastic region of the curves.

### 4.3 Animal Care and Surgical Procedures

All procedures described in this study were approved and performed in accordance with standards set out by the University of Ottawa Animal Care and Veterinary Services ethical review committee. Results are also reported in accordance with ARRIVE guidelines. Female Sprague Dawley rats ranging in weight from 250-300 grams were purchased from Charles River. The rats were anesthetized with isoflurane USP-PPC (Pharmaceutical partners of Canada) and injected subcutaneously with normal saline (Baxter) and enrofloxacin (Baytril). Laminectomies were performed at the T8-T9 level to expose the spinal cord. The dura was incised with micro scissors to expose the spinal cord. A hook was passed ventrally to ensure the entire cord was within the bend of the hook. The spinal cord was carefully lifted with the hook, and the whole cord was then cut with micro scissors. Both stumps of the spinal cord were carefully examined to confirm complete transection of the spinal cord and spinal roots at that level. Surgifoam 1972 (Ethicon) was inserted into the gap between the two spinal cord stumps. After several minutes, when hemostatic control was established, the surgifoam was removed, and the resulting gap size was measured. Before the surgery, animals were randomly assigned as a biomaterial or negative control. For animals assigned to the biomaterial group, a biomaterial scaffold was selected that best matched the gap distance of the stumps (range was typically 1 to 3.5 mm). A volume of 0.2 mL Fibrin glue (Tisseel) was then applied to the dorsal surface to stabilize the biomaterial. Negative control animals had 0.2 mL fibrin glue placed between the two stumps. The muscle layers were then reapproximated with 3-0 Vicryl sutures (Johnson & Johnson) while the skin was closed with Michel clips (Fine Science Tools). Following the surgery, rats had their bladders expressed manually three times a day and were monitored for any symptoms of weight loss, dehydration, pain and urinary infections.

### 4.4 Functional studies

The locomotor function of the rats was assessed weekly based on the Basso Beattie Bresnahan (BBB) open-field assessment^63^. The rats were placed in a 1-meter diameter arena every week and recorded for the entire duration of the study. At every time point at least 3 animals per condition were tested (Supplemental Table 1). Each of the five minutes videos of the rats was then scored by three blind observers and. Any substantial spasticity and reflexes/twitches in any of the four joints were ignored and confirmed with repeated views of the videos.

### 4.5. Retrograde Tracer Surgery

Rats with biomaterial and controls were randomly selected at 14 weeks post-injury to undergo retrograde tracer studies. Animals were anesthetized and maintained with isoflurane USP-PPC (Pharmaceutical partners of Canada). The retrograde tracer surgery and loading techniques were optimized from previous reports^27,75^. A laminectomy was then performed at the T13 level with a complete spinal cord transection performed as described in the previous spinal cord transections. In the spinal cord transection gap, a pledgets of surgifoam (5 mm^2^) were punched out and soaked in 4% FluoroGold (Fluorochrome) in saline. The FG soaked surgifoam was placed onto the rostral end of the spinal cord stumps. Petroleum jelly (Sherwood Medical) was then added to stabilize the surgifoam into place and prevent nonspecific labelling before the muscles were then closed with 3-0 Vicryl. The skin was then closed with Michel suture clips (Fine Science Tools). The rats were allowed to recover and were monitored for 7 days to allow the FG to travel across the scaffold. Axial sections rostral and caudal (n=7 for scaffold and n=3 for control) 2-3 mm from their respective scaffold position were sectioned and scanned. The remaining stubs of tissue were then repositioned and sectioned in an sagittal orientation. Sections were mounted with in VectaShield Vibrance (Vector Laboratories) before adding a coverslip. prior to scanning. For dapi-labelled FG slides, sections were incubated with 1ug/mL of DAPI (Thermo Scientific) prior to mounting. Negative controls were to test the unspecific wicking capacity of FG in the scaffold. Non-implanted scaffold material were incubated with 4% FG for 2 hours. Negative controls scaffolds were sectioned and imaged as the implanted scaffolds. Animals receiving FG retrograde had the scaffold dissected and had the rostral and caudal spinal cord positioned into blocks for analysis. The CNS tissue rostral and caudal of the scaffold were sequentially sectioned in the axial orientation from either ends of scaffold block. FG slides samples were imaged with a Nikon Ni-U Ratiometric Fluorescence Microscope with a 340/380 filter set. FG cell bodies were manually counted from axial sections. All image processing was carried out with Nikon Elements software (Nikon).

### 4.6 Spinal Cord and Biomaterial Resection

For harvesting of the tissue, animals were sacrificed by intraperitoneal injection of 0.7–1.0 ml of sodium pentobarbital (65 mg/ml) and underwent intracardiac perfusion with 500 mL of normal saline and 0.5 U/mL heparin solution. The rats were then perfused with 500 mL of 4% paraformaldehyde (Sigma-Aldrich) in 0.1mM PBS(Sigma-Aldrich). The entire spinal cord and brain were then dissected out and further fixed overnight in 4% paraformaldehyde and 0.1mM phosphate buffer solution at 4°C. The tissue was then removed from the fixation solution and incubated in 30% sucrose (Sigma-Aldrich) 1% sodium azide (Sigma-Aldrich) in PBS for 24 hours at 4°C. The biomaterial and the surrounding tissue were then frozen and mounted in Optimum Cutting Temperature compound (Stephens Scientific), with the remaining tissue fixed into paraffin. For tracer animals, the entire spinal cord and brain were frozen in individual blocks. Axial and sagittal sections of the tissue at 7–10 μm thick were then processed and mounted onto cold Superfrost Plus slides (Fisher Scientific) by the PALM Histology Core Facility at the University of Ottawa. Slides were stored in a -80°C freezer until stained and mounted.

### 4.7 Histology and Immunohistochemistry

Frozen sections and paraffin section were stained with hematoxylin-eosin (H&E) and Masson Trichrome (MT). Frozen sections were completely dried overnight and rehydrated in 1X TBST buffer and blocked for 30 minutes with Rodent Block R (Biocare). Sections were then incubated overnight at 4°C with the following primary antibodies rabbit AB5804 GFAP (1:2000, Millipore) and mouse N0142 NF200 (1:3000, Sigma). The following day, the sections were washed with 1X TBST and then incubated with secondary antibodies: AB150077 Goat anti-rabbit 568 (Abcam) or AB175473 Goat anti-mouse 568 (Abcam) at 1:500 dilution for 2 hours in the dark at room temperature. For Rabbit 5HT S5545 (Sigma), Rabbit S-100 S2644 (Sigma), Rabbit Laminin L9393 (Sigma); frozen sections were dried overnight and fixed in a methanol/acetone solution for 10 minutes and allowed to dry at room temperature for an hour. The slides were then rehydrated with 1x TBST buffer and treated with Rodent Block RBR962G (Biocare) Sections were then incubated for 90 minutes at room temperature with the following primary antibodies concentrations: Rabbit 5HT (1:1500), Rabbit S-100 (1:250), Rabbit Laminin (1:25). The sections were washed with 1XTBST prior to incubation with the secondary (1:500) Donkey anti-rabbit 568 (Abcam) for 2 hours in the dark at room temperature. All sections were washed with 1X TBST, incubated with 1ug/mL of DAPI (Thermo Scientific), washed, and then mounted in VectaShield Vibrance (Vector Laboratories) before adding a coverslip. The slides were kept in the dark at -20°C for a maximum of a week prior to imaging. A third-party pathologist reviewed H&E slides to provide interpretation. CD31 IHC staining was performed on formalin fixed paraffin embedded tissue sections using the Leica Bond(tm) system using a modification of protocol F that eliminates the post primary step when using rabbit antibodies on rat tissue. Sections stained with Rabbit anti-CD31 (Novus Biologicals) were pre-treated using heat mediated antigen retrieval with a Sodium Citrate buffer (pH 6, epitope retrieval solution 1) for 20 minutes. The sections were then incubated using a 1:100 for 30 minutes at room temperature and detected using an HRP conjugated compact polymer system. Slides were then stained using DAB as the chromogen, counterstained with Hematoxylin, mounted and cover slipped.

### 4.8 Microscopy

Micrographs of colorimetric stains were captured using Zeiss AxioScan Z1 slide Scanner (Zeiss) equipped with 10x objective and analyzed using ZenBlue (Zeiss, Canada) and ImageJ software. Phase microscopy was carried out on an A1R TiE inverted optical microscope (Nikon). Fluorescence imaging of tissue sections stained with NF200 and GFAP antibodies was carried out on a Nikon A1RsiMP Confocal Workstation with a 32-detector array for spectral imaging with 6 nm resolution detectors (Nikon). As lignocellulose can be autofluorescent, spectral linear unmixing was achieved by obtaining autofluorescence spectra of unstained scaffolds (Supplementary Fig. 18). The spectral profile of the 568 antibody was then unmixed from any scaffold autofluorescence, and the resulting unmixed z-stack was 3D deconvolved to create a projection. Although antibodies have the same 568nm wavelength fluorophore, for clarity, the signal is presented in false-colour to distinguish from one another.

We also sought to determine the area of GFAP signal, normalized by the total tissue area. In these cases, saggital sections of animals receiving the scaffold (n=7) and control (n=4) were stained with GFAP (red) and DAPI. The rostral/caudal interfaces of the scaffold with several mm of cord tissue was scanned. We analyzed a total of 26 sections for scaffolds and 19 for control conditions. For animals receiving the scaffold, the interface between the scaffold and the host was defined as the boundary. Regions of interest (ROI) of 500 × 1500 microns were defined along the scaffold interface with the long axis reaching into the spinal cord tissue The total “tissue area” was determined from the outer boundary formed by the densely packed DAPI positive nuclei. Although in an ideal setting tissue would be perfectly sectioned with no tearing or lift-off, utilizing the DAPI signal allowed us to ensure a fair comparison in the very small number of cases where the tissue integrity was less than perfect. The scar area was then determined by thresholding and the GFAP signal, followed by area quantification using the particle analyzer plugin in Fiji. The normalized area was calculated by dividing the GFAP area by the tissue area. The GFAP scar area was identified according to the morphology of the GFAP labelled cells, which clearly delineated the ischemic core from the ischemic penumbra. This procedure was also carried out for control tissues where the interface was easily identified as the edge of spinal cord stump and the cyst. Normalized areas were then expressed as mean ± S.E.M.

### 4.9 Scanning Electron Microscopy

The structure of cellulose was studied using SEM. Globally, scaffolds were fixed in an electron microscopy grade 4% PFA (Fisher Scientific) and dehydrated through successive gradients of ethanol (50%, 70%, 95% and 100%). The biomaterial was then dried with a critical point dryer (CPD) (SAMDRI-PVT-3D). Samples were then gold-coated at a current of 15mA for 3 minutes with a Hitachi E-1010 ion sputter device. SEM imaging was conducted at voltages ranging from 2.00–10.0 kV on a JSM-7500F Field Emission SEM (JEOL).

### 4.10 Statistics

All values reported here are the average ± standard error of the mean. Statistical analyses of the BBB scoring and scaffold volume were performed with one-way ANOVA by using SigmaStat 3.5 software (Dundas Software Ltd). A value of p<0.05 was considered statistically significant.

## ACKNOWLEDGMENTS

We would like to thank Dr. Holly Orlando and the veterinary technicians of the Animal Care and Veterinary Service team of the University of Ottawa; Anne-Renée Desjardins, Catherine Thibault, Roxanne Cote, Caroline Côté, Anik Baillot, Amanda Wells, Melissa Washington, Christine Kitchen, Pascale Beaudry, Catherine Lépine Bisson, and Tami Janveau.The authors would like to acknowledge the technical support provided by Louise Pelletier Histology Core Facility, Department of Pathology and Laboratory Medicine, University of Ottawa also like to thank Dr. Ana Giassi, Dr. Sharlene Faulkes, Dr. Li Dong, Eric Labelle and Dr. Ziba Jaberansari for their help with histological processing. We would like to thank Dr. Rafay Axhar and Dr. John Woulfe, for their histopathological analysis. We thank Andrew Ochalski, Dr. Wissam Nakhle and Dr. Gregory Cron for their microscopy and image-processing assistance. The authors wish to thank Dr. Yun Liu for assistance with SEM imaging. We also thank Dr. Sebastian Hadjiantoniou and Dr. Sophie Chagnon-Lessard for surgical assistance. This work was supported in part by grants to AEP from the Natural Sciences and Engineering Research Council Discovery Grant, a Canada Research Chair, the Canada Foundation for Innovation, the Li Ka Shing Foundation and the University of Ottawa.

## DATA AVAILABILITY

The research data required to reproduce these findings is available upon reasonable request from the corresponding author. The study is reported in accordance with ARRIVE guidelines.

## AUTHORSHIP CONTRIBUTION STATEMENT

**Daniel J. Modulevsky:** Conceptualization; Data curation; Formal analysis; Investigation; Methodology; Roles/Writing - original draft; Writing - review & editing; Visualization; Supervision. **Charles M. Cuerrier** Data curation; Formal analysis; Investigation; Methodology; Supervision; Writing - Review & Editing, Funding acquisition. **Maxime Leblanc-Latour:** Data curation; Investigation, Writing - Review & Editing, Methodology. **Ryan J. Hickey:** Data curation; Investigation, Writing - Review & Editing, Methodology. **Isabel Shores:** Data curation; Investigation, Writing - Review & Editing, Methodology. **Ras-Jeevan K. Obhi:** Data curation; Investigation, Writing - Review & Editing, Methodology. **Ahmad Galuta:** Data curation; Investigation, Writing - Review & Editing, Methodology. **Krystal L. A. Walker:** Data curation; Investigation, Writing - Review & Editing, Methodology. **Eve C. Tsai:** Conceptualization; Formal analysis; Investigation; Methodology; Roles/Writing - original draft; Writing - review & editing; Resources; Visualization; Supervision. **Andrew E. Pelling:** Conceptualization; Formal analysis; Investigation; Methodology; Roles/Writing - original draft; Writing - review & editing; Resources; Data Curation; Visualization; Supervision; Project administration; Funding acquisition

