## supplemental files for "Plant Scaffolds Support Motor Recovery and Regeneration in Rats after Traumatic Spinal Cord Injury"

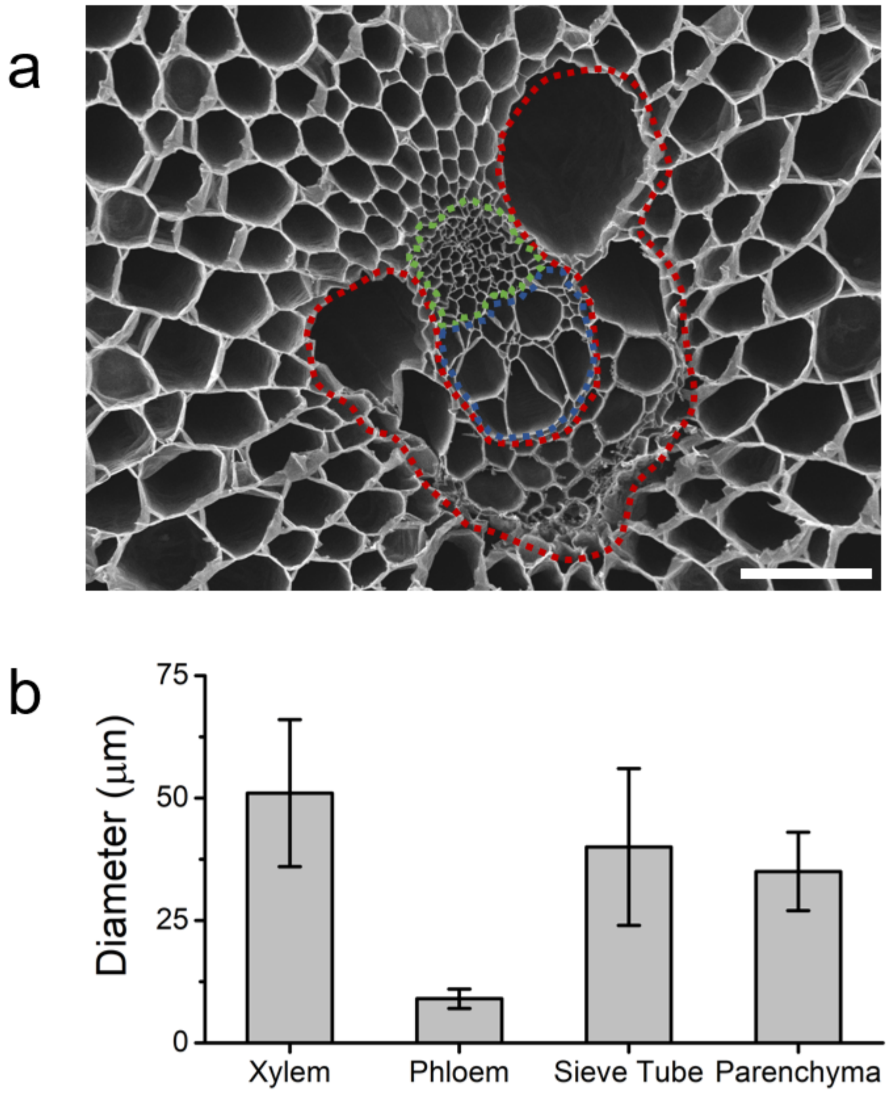


**Supplementary Fig. 1 Different structures of the vascular bundle. a)** A scanning electron microscope image of the surface of a decellularized scaffold revealing a single vascular bundle (VB) and surrounding parenchyma tissue (bar=100µm). The elements of the VB are highlighted to show the distinct channel architectures. The xylem (red) are channels that run the entire length of the asparagus and transport water within the plant. The phloem (blue) transport sugars within the asparagus from photosynthetic cells to non-photosynthetic cells. Phloem structures differ from xylem as they contain highly perforated sieve elements along their length. The sieve tubes are outlined in blue. The sieve tubes contain specialized cells with no nucleus that have roles in transporting carbohydrates/ messaging molecules throughout the plant. **b)** Characteristic diameters of the various elements of the vascular bundle, xylem channels (51±15µm), sieve tubes (40±16µm), parenchyma (35±8µm) and the phloem (9±2µm).

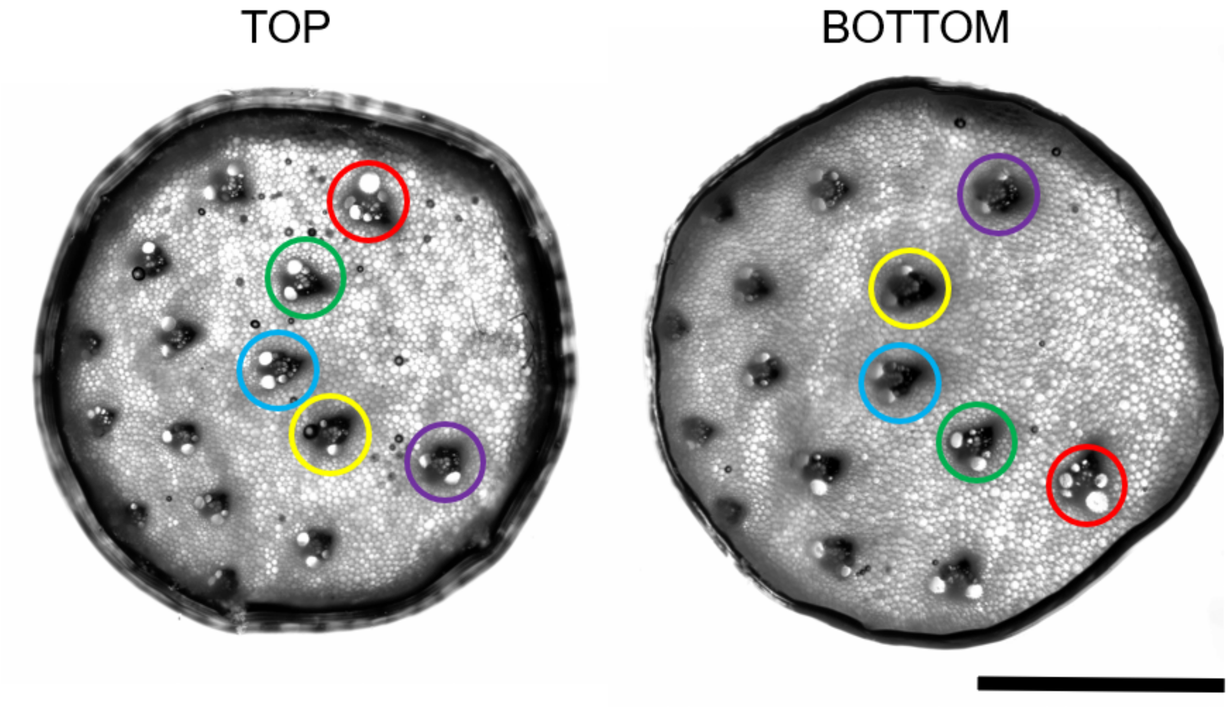


**Supplemental Fig 2. The two opposite ends of the scaffold.** Phase contrast image of the entire surface of the scaffolds revealing the distribution of the VB within the scaffold on the top surface and the emergence of the same VB in the nearly the exact position in the bottom surface (bar=2mm).

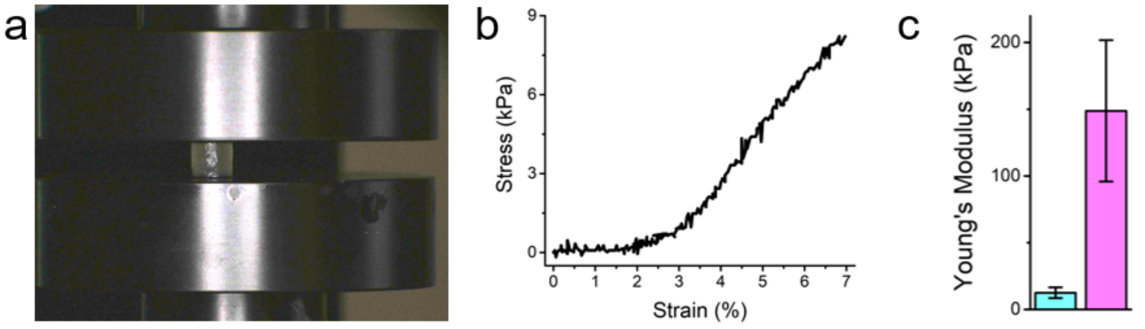


**Supplemental Fig 3. The young modulus of the SCI scaffold prior to implantation. a)** The SCI scaffold loaded into the CellScale UniVert compression platform. **b)** The elastic deformation of the stress strain curve fort the SCI scaffold used to determine the scaffold. **c)** The quantified young's modulus of the SCI scaffold along the parallel axis (blue) and perpendicular to the long axis (pink).

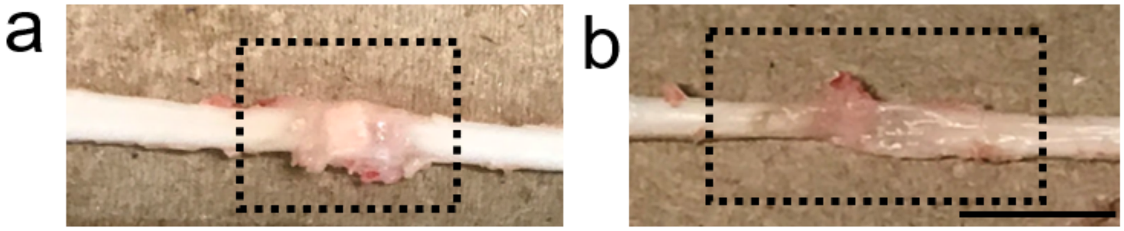


**Supplemental Fig 4. Dissection of entire CNS tissue of animals receiving the scaffolds and control after 28 weeks. a)** During the dissection, the scaffold was well integrated and could not be removed from the two stumps of the spinal cord. The scaffold was so well adhered that the it could support the weight of the CNS when lifted. **b)** The control groups were difficult to remove as the stumps loosely adhered via scar and connective tissues. The damage to the surrounding spinal cord tissue can also be observed much further in both the rostral and caudal spinal cord stumps (bar=1cm).

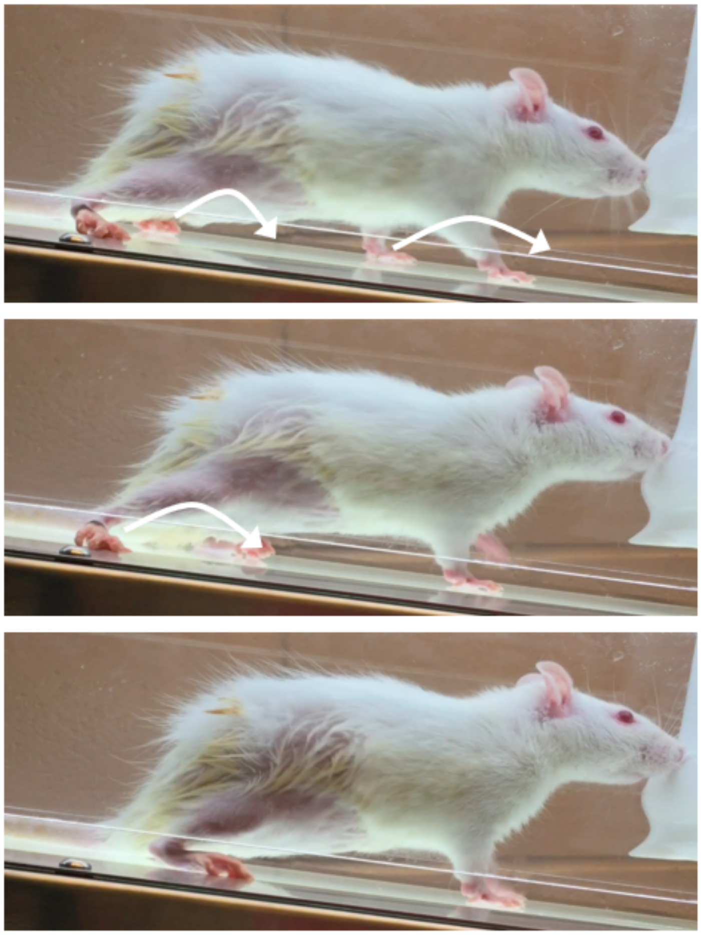

**Supplemental Fig 5. Coordinated hind and forelimb load bearing steps in biomaterial treated rats.** Three animals receiving the scaffold, that demonstrated increased motor recovery and were placed onto a treadmill for gait analysis. On the treadmill, an animal exhibited select coordinated load bearing steps.

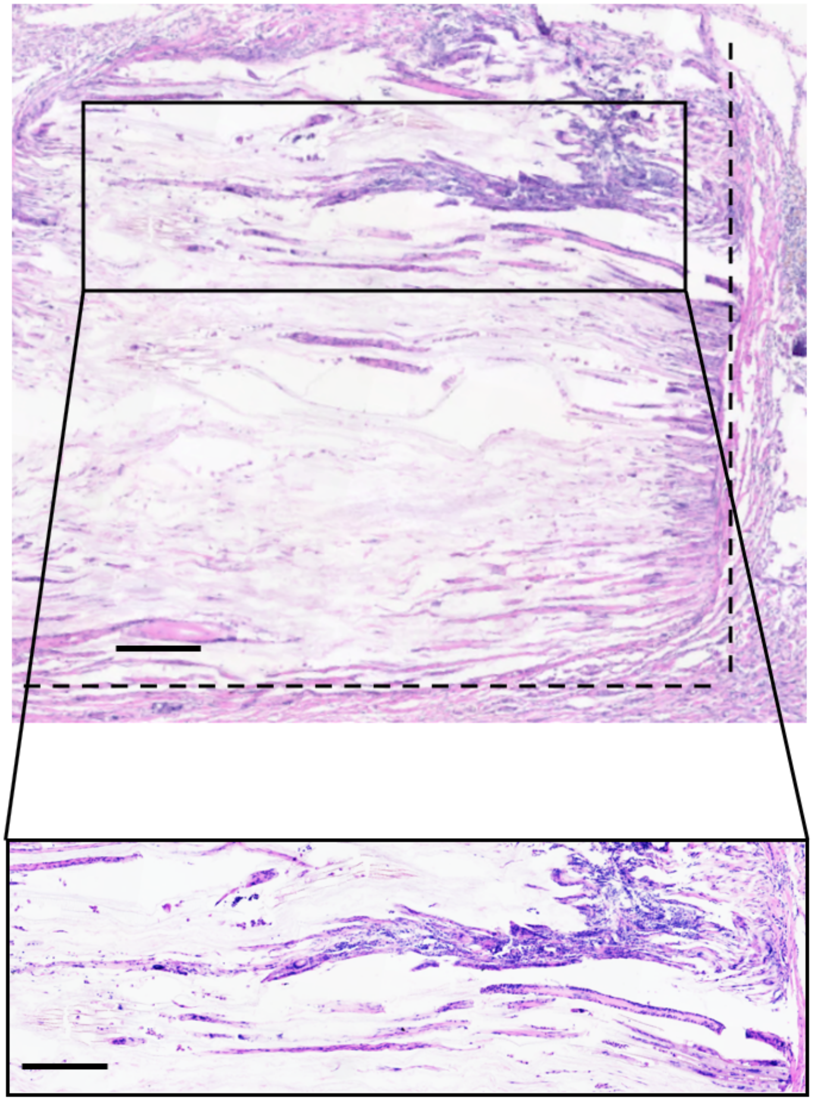

**Supplemental Fig 6. Hematoxylin and Eosin staining after twenty-eight weeks of implantation. a)** Sagittal macro view of H&E staining of SCI scaffold at the T8-T9 vertebrate after twenty eight weeks of implantation. The vascular bundles can be seen throughout the scaffold infiltrated with host cells. The majority of the infiltrating host cells are confined to the vascular bundles however the parenchyma tissue is not void of cells (bar=200µm). **b)** Sagittal H&E staining of SCI scaffold at the T8-T9 vertebrate after 28 weeks post SCI injury. The vascular bundles can be seen completely infiltrated the entire length of the scaffold from the two opposing stumps. As with the coronal H/E there is very little foreign body response to the scaffolds. Even after 28 weeks there is no degradation to the scaffold and the dimensions of the scaffold are still apparent (represented in dashed lines) and have withstood any internal pressure (bar=200µm).

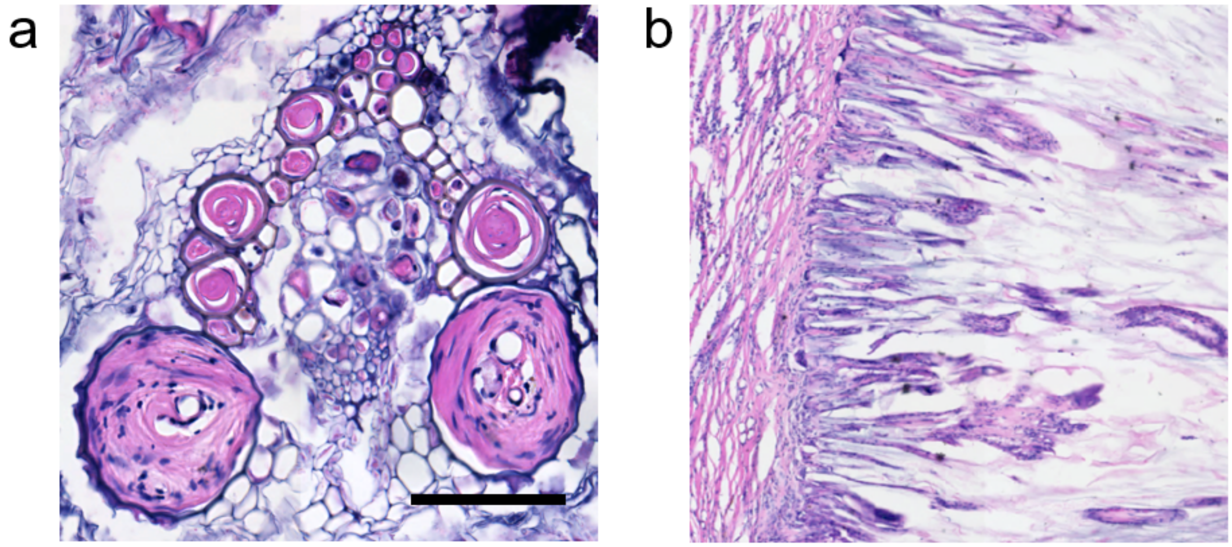


**Supplemental Fig 7. Hematoxylin and Eosin staining after 28 weeks of implantation. a)** Axial view of H&E staining of SCI scaffold at the T8-T9 vertebrate after 28 weeks of implantation. The vascular bundles can be seen throughout the scaffold infiltrated with host cells. The majority of the infiltrating host cells are confined to the vascular bundles however the parenchyma tissue is not void of cells (bar=200µm). **b)** Sagittal H&E staining of SCI scaffold at the T8-T9 vertebrate after fourteen weeks post SCI injury. The vascular bundles can be seen completely infiltrated the entire length of the scaffold from the two opposing stumps. As with the coronal H&E there is very low foreign body response to the scaffolds. Even after 28 weeks there is no degradation to the scaffold and the dimensions of the scaffold are still apparent (represented in dashed lines) and have withstood any internal pressure (bar=200µm).

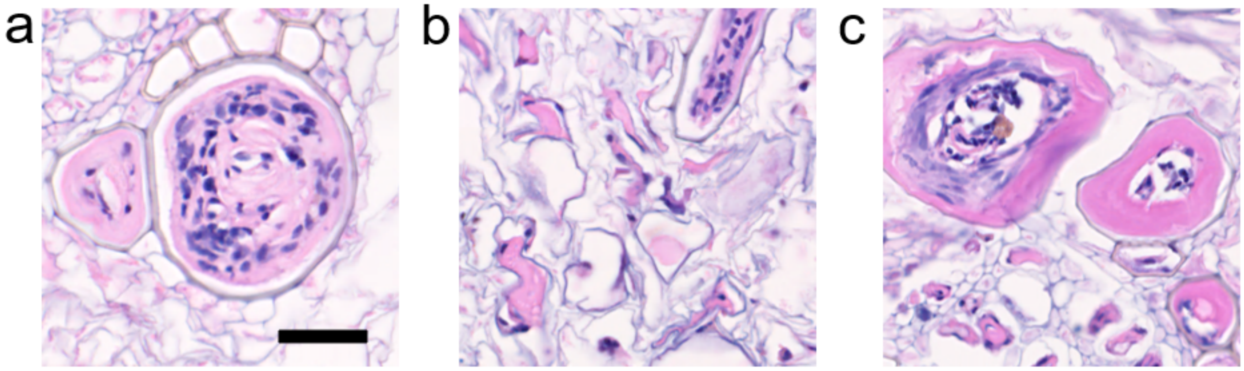


**Supplemental Fig 8. High magnifications of Hematoxylin and Eosin demonstrating specific structures. a)** Mammalian granulation scar tissue can be seen in the xylem channels of the VB. This scar tissue that typical prevents the extension of axons are seen sequestered in the xylem channel leaving the rest of the surrounding parenchyma tissue bar. **b)** The spindle shaped nucleus of active fibroblast can be seen in the surrounding parenchyma tissue. **c)** Blood vessels with thick endothelial linings can be observed throughout the VB and in the parenchyma tissue supplying the host cells (bar=50µm for all).

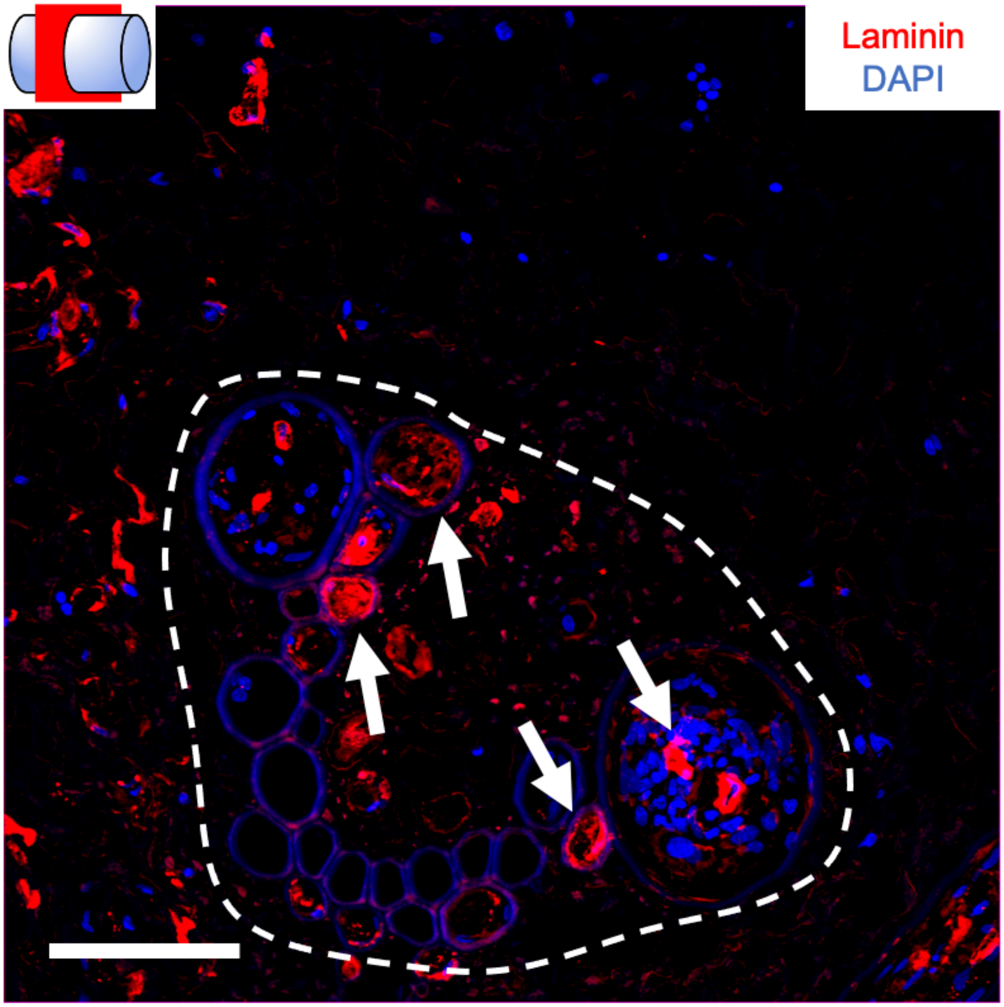


**Supplemental Fig 9. Laminin deposition with the scaffold after 28 weeks.** Axial section within the scaffold of laminin deposition (red) and DAPI (blue) at 28 weeks (bar=100µm). Laminin rich blood vessels can be seen within the larger channels of the VB. The VB is within the dotted line and arrows indicate some of the laminin-rich structures. Laminin can also be seen throughout parenchyma outside of the VB.

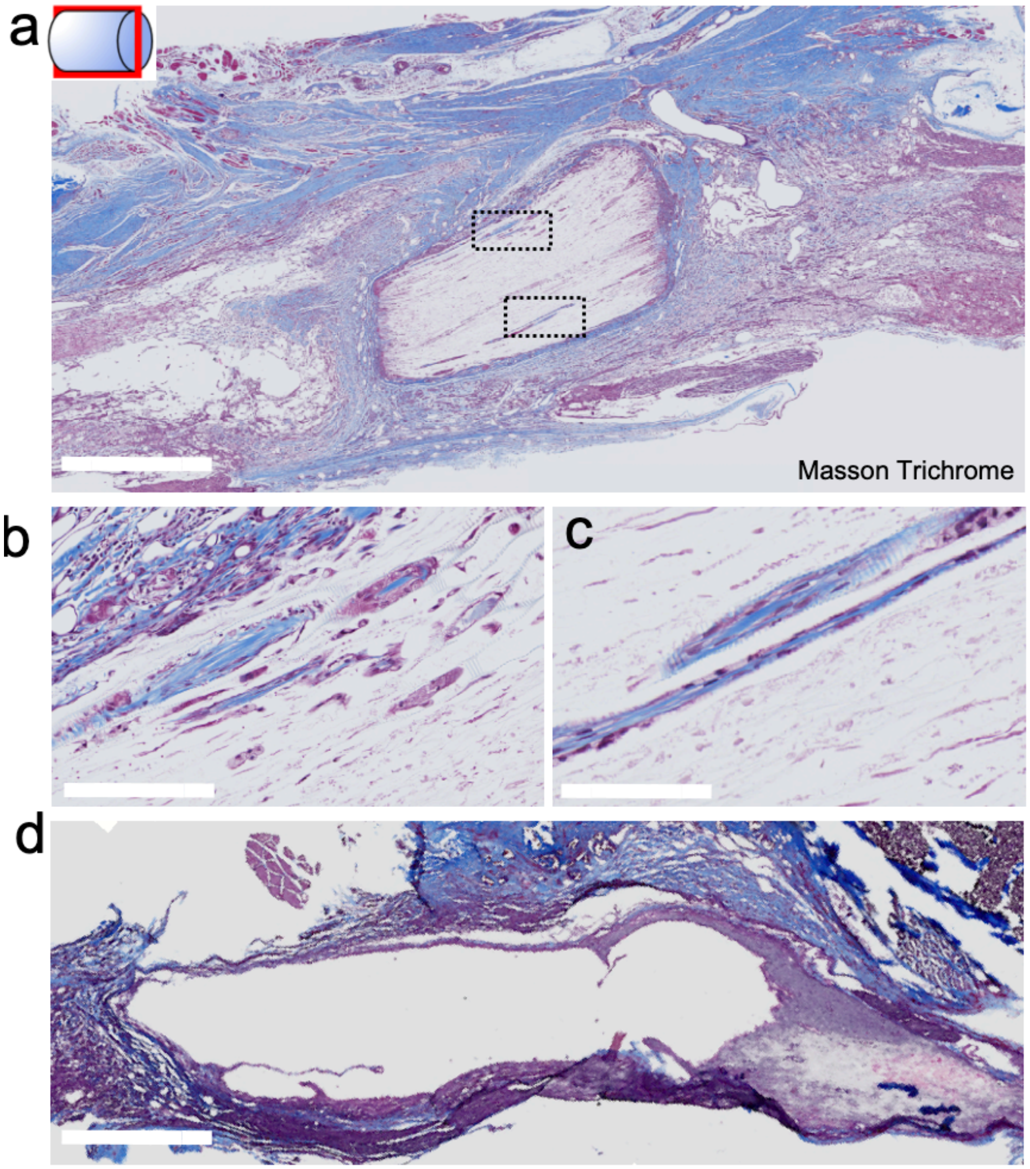


**Supplemental Fig 10. Sagittal Masson Trichrome stain of the injury site after 14 weeks.** The scaffold is observed integrated within the initial injury site of the spine. Host tissue is observed surrounding the scaffold and within the VBs and the parenchyma even at the midpoint of the scaffold. Both stumps appear to be stabilized by the presence of the scaffold which may also provide mechanical support. The stumps appear adhered with host tissue growing throughout the scaffold and on the surfaces (bar =1 mm). **b) and c)** Higher magnification of the boxed ROIs in **a)**. Cells and collagen deposition from both stumps and can be observed within the VB channels and not the surrounding parenchyma (bar=100μm). This is consistent with laminin staining. **d**) Sagittal Masson Trichrome stain of the surrounding spinal cord injury site animals not receiving the scaffold after 14 weeks (bar=1mm).

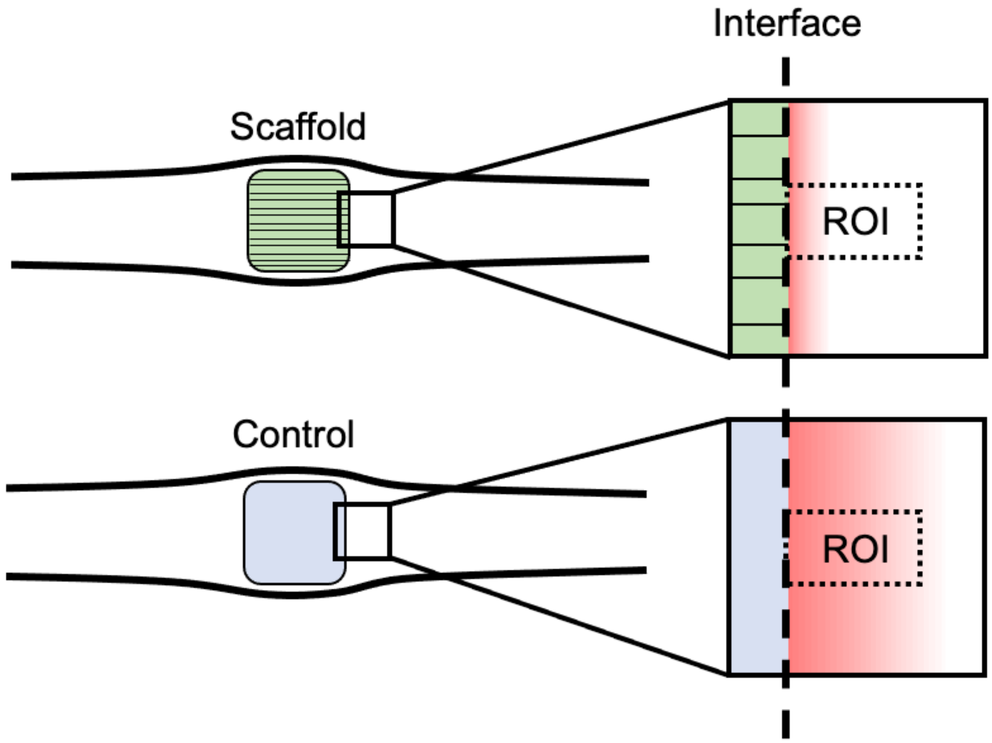


**Supplemental Fig 11. Illustration of the procedure to determine normalized GFAP area.** Sagittal sections of animals receiving the scaffold (n=7) and control (n=4 stained with GFAP (red) and DAPI were analyzed. The rostral/caudal interfaces of the scaffold with several mm of cord tissue was scanned. We analyzed a total of 26 sections for scaffolds and 19 for control conditions. For animals receiving the scaffold, the interface between the scaffold and the host was defined as the boundary. Regions of interest (ROI) of 500 x 1500 microns were defined along the scaffold interface with the long axis reaching into the spinal cord tissue. The total "tissue area" was determined from the outer boundary formed by the densely packed DAPI positive nuclei. Although in an ideal setting tissue would be perfectly sectioned with no tearing or lift-off, utilizing the DAPI signal allowed us to ensure a fair comparison in the very small number of cases where the tissue integrity was less than perfect. The scar area was then determined by thresholding and the GFAP signal, followed by area quantification using the particle analyzer plugin in Fiji. The normalized area was calculated by dividing the GFAP area by the tissue area. The GFAP scar area was identified according to the morphology of the GFAP labelled cells, which clearly delineated the ischemic core from the ischemic penumbra. This procedure was also carried out for control tissues where the interface was easily identified as the edge of spinal cord stump and the cyst. Normalized areas were then expressed as mean ± S.E.M.

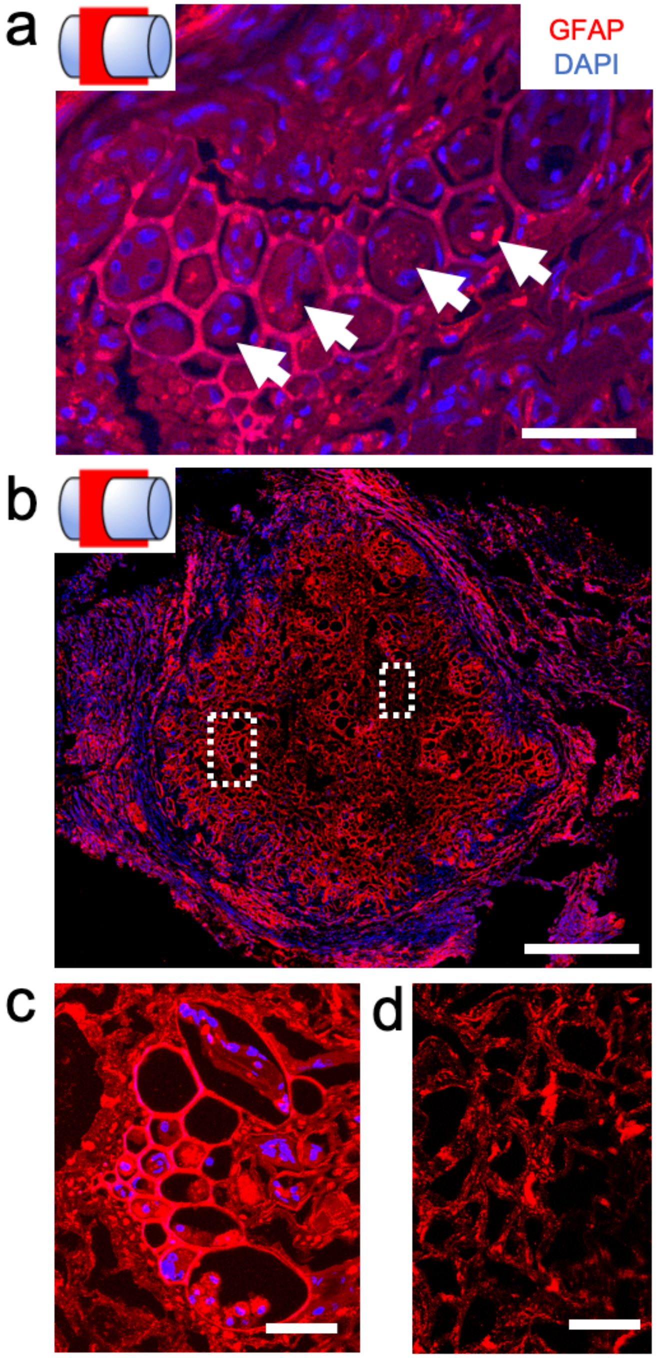


**Supplemental Fig 12. Axial section of GFAP staining within the scaffold at 14 weeks. a)** VBs with the larges to the channel elements infiltrated with GFAP labelled tissue (bar = 50 microns) **b)** Axial section global view of the GFAP labelled scaffold (bar = 500 microns). **c)** Magnified ROI of a VB with GFAP signal within the channels (bar= 50 microns for c and d). **d)** Magnified ROI of parenchyma with reduced GFAP labelling.

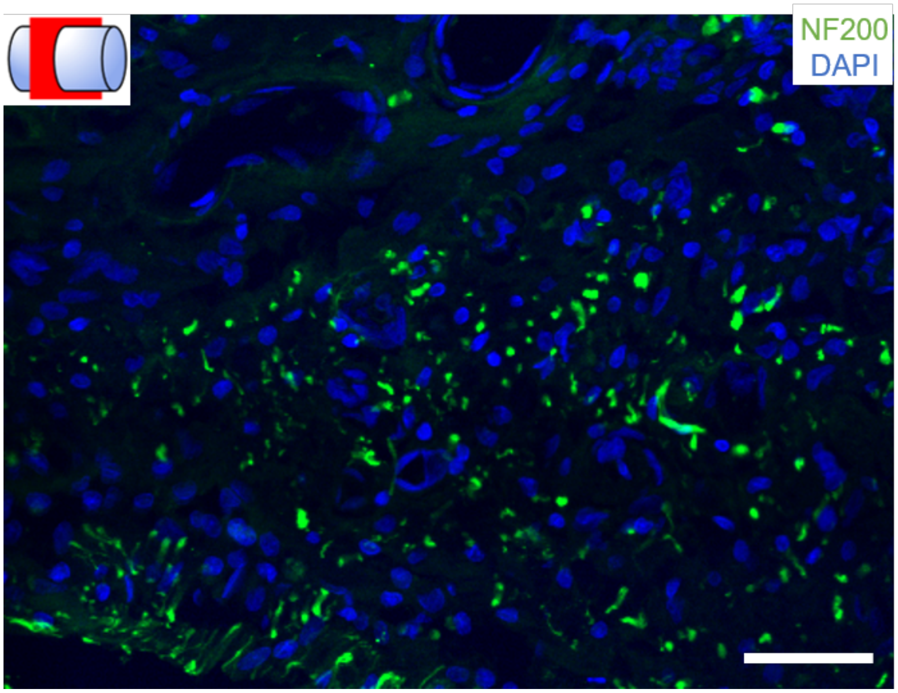


**Supplemental Fig 13. NF200 labelled axons within the rostral stump of a control animal after 14 weeks of implantation.** Animals not receiving the scaffold were sectioned in the axial orientation. In control animals, NF200 labelled axons were only observed in the scar tissue of the stumps (bar=20µm).

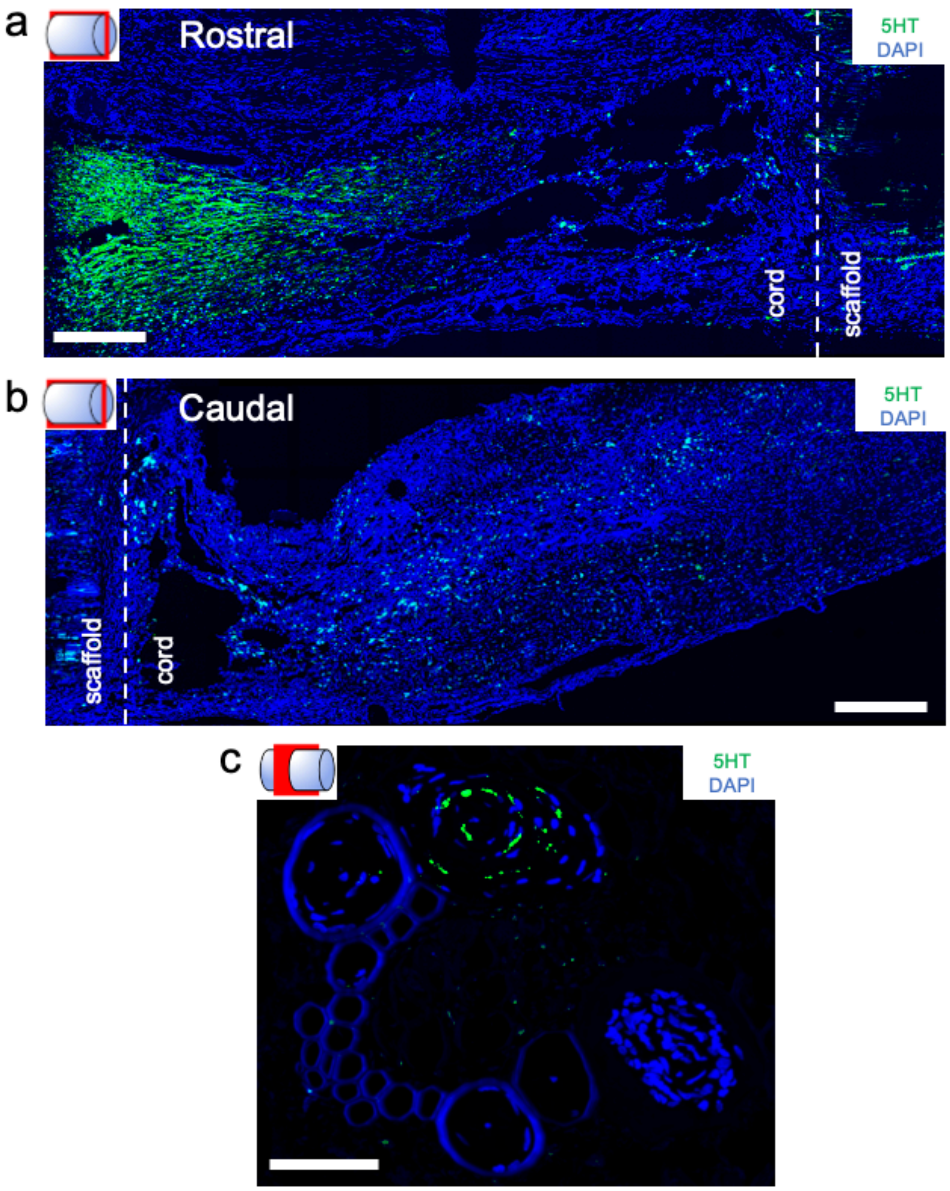


**Supplemental Fig 14. Serotonergic axons stained with 5HT in animals receiving scaffolds (n=4) and control animals (n=3). a)** Rostral end, sagittal view of the spinal cord-scaffold interface (dotted line) stained for 5HT (green) and DAPI (blue) (bar=500μm, for **a** and **b**). As expected, a number of 5HT positive axons appear to have sprouted rostral to the injury site. The data is consistent with control animals. **b)** Caudal end, sagittal view of spinal cord-scaffold interface (dotted line). In this case some point-like 5HT positive signals do appear in the tissue but the density is low compared to the rostral cord. some potential structures. **c)** Axial section of 5HT axons and DAPI within the scaffold. 5HT-positive labelling in axons was observed in the vicinity of the VBs at both 14 and 28 weeks (bar=80μm). However, their density was very low and they were only observed in a limited number of locations throughout the entire scaffold. In general, 5HT serotonergic neurons were identified rostral to the injury/scaffold but they do not appear to descend to the caudal region of the cord in great number/density.

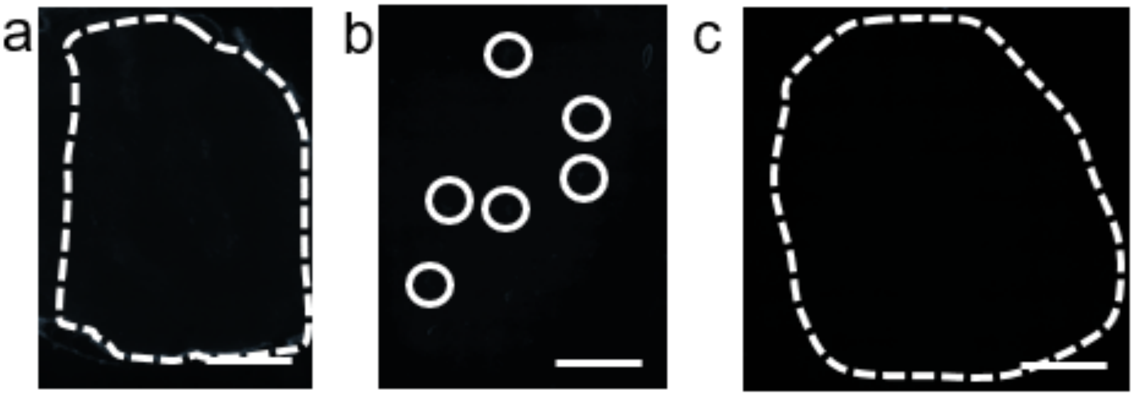


**Supplemental Fig 15. Fluoro-Gold retrograde axonal tracing control experiments 14 weeks post injury. a)** Axial spinal cord sections of the rostral end of the transection in a 14-week animal that did not receive a scaffold. There is a lack of FG signal throughout the spinal cord cross section with only limited nonspecific signal observed in the dura of the spinal cord (bar=500µm). The tissue is outlined with a white dotted line. **b)** Axial section of a non-implanted scaffold material directly loaded with 4% FG in a manner similar to the procedure performed at the secondary transection site. The FG does not adhere to cellulose or lignocellulose materials of the scaffold leading to a lack of signal (bar=1mm). Individual VBs are circled in white. **c)** Axial spinal cord section of an animal that did not receive the FG stain (bar=500µm). All data presented here was acquired and processed with the same settings and parameters as the data presented in Fig. 4. The images primarily appear black due to the lack of signal as expected for these control experiments. The tissue is outlined with a white dotted line.

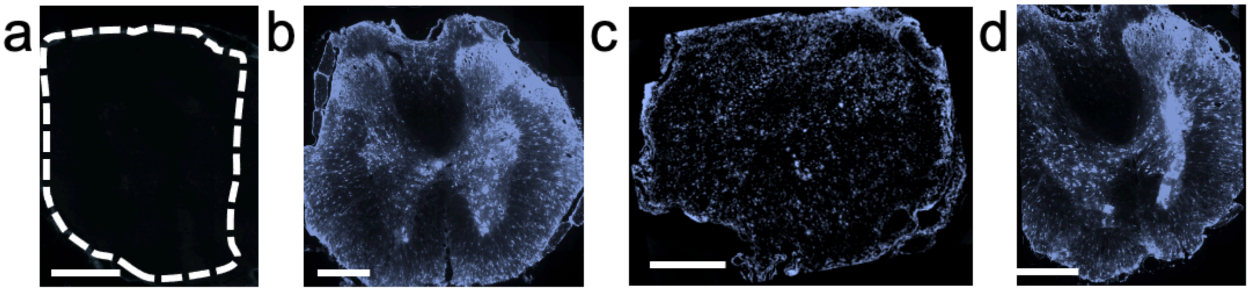


**Supplemental Fig. 16 Fluoro-Gold (FG) retrograde axonal tracing in the rostral and caudal ends of the spinal cord 14 weeks post injury. a)** Axial section of the spinal cord rostral to the injury site in an animal that did not receive scaffold. This is the same image from Supplementary Fig. 7a. Due to the lack of FG transport across the injury site, any observable signal is very low. Due to this, the tissue is outlined with a white dotted line to guide the reader (bar=500μm). **b)** Axial section of the spinal cord caudal to the injury site in an animal that did not receive scaffold. Unlike in **a**, neuron cell bodies labelled with FG are visible in the dorsal and ventral horns of the spinal cord as expected (bar=500μm). **c)** Axial section of the spinal cord rostral to the injury site in an animal that did receive a scaffold. Due to FG transport across the scaffold via projecting axons, some signal is observed in this part of the cord (bar=500μm). Interestingly, the diffuse pattern and distribution of FG positive axons qualitatively matches the distribution of NF200 positive axons that were imaged directly in the scaffold itself (Fig. 3). **d)** Axial section of the spinal cord caudal to the injury site in an animal that did receive a scaffold, which appears very similar to **b** as expected (bar=500μm). The results reveal that while axons can project through the scaffold, they do so in a manner that does not preserve the global morphology and architecture of the healthy tissue. However, this does not come at a surprise given the extensive damage and trauma at the injury site.

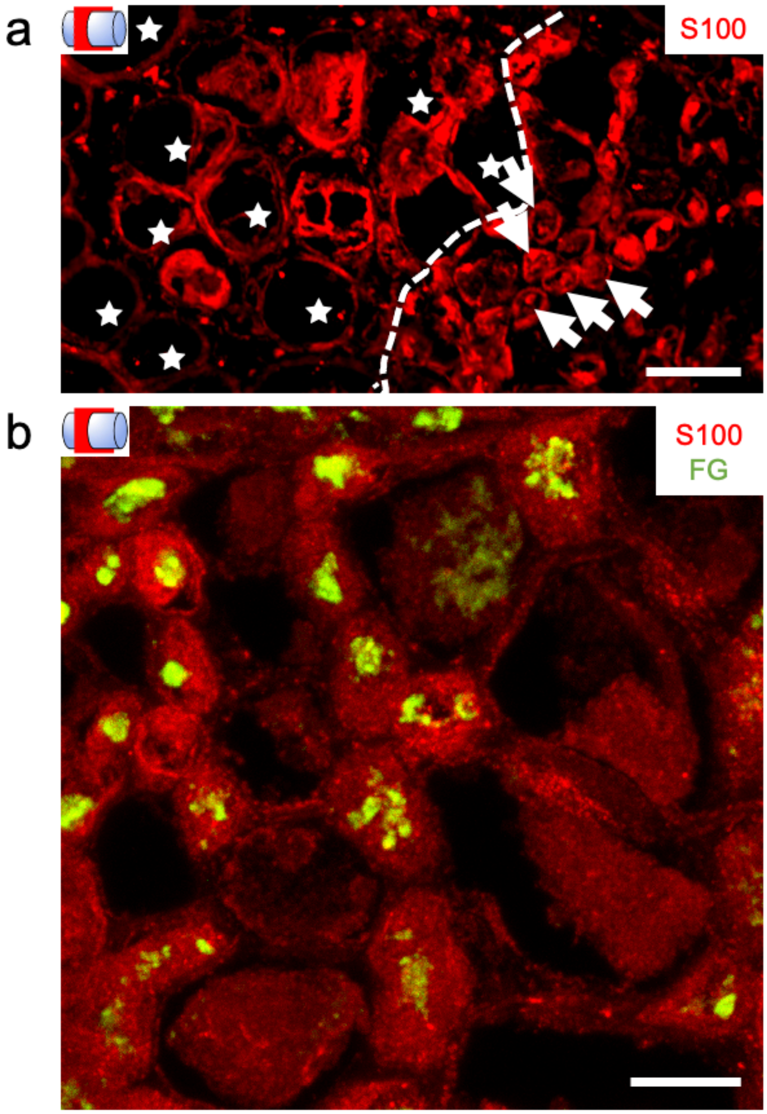


**Supplemental Fig 17. S100 labelled Schwann cell infiltration within the scaffold after 28 weeks. a)** Axial section within the scaffold of S100 labelled Schwann cells scaffold (red) migrating into the at 28 weeks (bar=50µm). The Schwann cells are within the smaller elements of the parenchyma (arrows) rather the unobstructed channels of the VBs (★). **b)** An axial section within a scaffold from an animal which received the FG retrograde tracer at 28 weeks was also stained for S100. Interestingly, in the parenchyma some S100 positive structures were found to be colocalized with FG (bar=10µm).

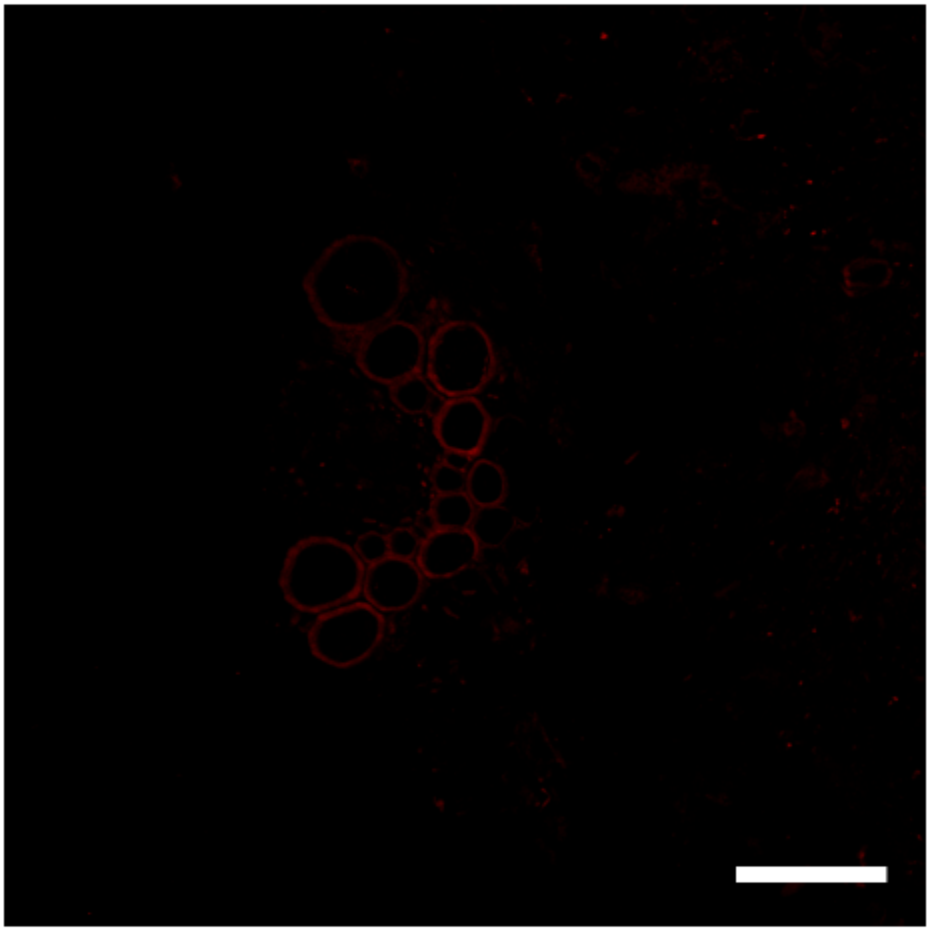

**Supplemental Fig. 18. Negative Control for Secondary GAR and GAM 568 antibody**
**a)** Coronal section of a 28-week implanted scaffold stained only with the secondary Goat anti-rabbit and Goat anti-mouse 568 antibody. The lack of signal demonstrates that the secondary antibodies did not display any non-specific binding with the cellulose scaffold or host native tissue. Background autofluorescence can be seen in the VB due to their high lignocellulosic content (bar=100 μm). In this study we utilized spectral unmixing to further remove autofluorescence based on its characteristic spectral signature.

**Supplemental Table 1.  Number of animals used in the BBB scale at each time point**

| Week | Scaffold | Control |
|---|---|---|
| **1** | 23 | 11 |
| **2** | 23 | 11 |
| **3** | 23 | 11 |
| **4** | 22 | 11 |
| **5** | 22 | 11 |
| **6** | 22 | 11 |
| **7** | 22 | 11 |
| **8** | 22 | 11 |
| **9** | 22 | 11 |
| **10** | 15 | 8 |
| **11** | 15 | 8 |
| **12** | 15 | 8 |
| **13** | 15 | 8 |
| **14** | 15 | 8 |
| **21** | 5 | 3 |
| **28** | 5 | 3 |
